# Multifaceted modes of γ-tubulin complex recruitment and microtubule nucleation at mitotic centrosomes

**DOI:** 10.1101/2022.09.23.509043

**Authors:** Zihan Zhu, Isabelle Becam, Corinne A. Tovey, Eugenie C. Yen, Fred Bernard, Antoine Guichet, Paul T. Conduit

## Abstract

Microtubule nucleation is mediated by γ-tubulin ring complexes (γ-TuRCs). In most eukaryotes, a GCP4/5/4/6 “core” complex promotes γ-tubulin small complex (γ-TuSC) association to generate cytosolic γ-TuRCs. Unlike γ-TuSCs, however, this core complex is non-essential in various species and absent from budding yeasts. In *Drosophila*, Spindle defective-2 (Spd-2) and Centrosomin (Cnn) redundantly recruit γ-tubulin complexes to mitotic centrosomes. Here we show that Spd-2 recruits γ-TuRCs formed via the GCP4/5/4/6 core, but that Cnn can recruit γ-TuSCs directly via its well-conserved CM1 domain, similar to its homologues in budding yeast. When centrosomes fail to recruit γ-tubulin complexes, they still nucleate microtubules via the TOG domain protein Mini-spindles (Msps), but these microtubules have different dynamic properties. Our data therefore help explain the dispensability of the GCP4/5/4/6 core and highlight the robustness of centrosomes as microtubule organising centres. They also suggest that the dynamic properties of microtubules are influenced by how they were nucleated.

## Introduction

During cell division, centrosomes act as major microtubule organising centres (MTOCs) to nucleate and organise microtubules that contribute to mitotic spindle formation (Conduit et al., 2015). Centrosomes comprise a pair of centrioles that recruit and are surrounded by the pericentriolar material (PCM). The PCM is a large collection of proteins and is the predominant site of microtubule nucleation and organisation during mitosis. On entry into mitosis, centrosomes expand their PCM in a process called centrosome maturation (Khodjakov and Rieder, 1999; Piehl et al., 2004). This is particularly dramatic in *Drosophila* cells because interphase centrosomes have very little PCM and do not organise microtubules, while mitotic centrosomes have relatively large amounts of PCM and robustly organise microtubules (Rogers et al., 2008). This makes *Drosophila* centrosomes ideal for the study of mitotic PCM assembly and microtubule nucleation.

γ-tubulin ring complexes (γ-TuRCs) are important PCM clients because they template the nascent assembly of microtubules (microtubule nucleation) (Tovey and Conduit, 2018; Farache et al., 2018; Kollman et al., 2011). Along with actin and Mozart proteins, they comprise a single-turn helical arrangement of 14 laterally associated “spokes”, each made from a γ-tubulin complex protein (GCP, or “Grip” protein in *Drosophila*) and a γ-tubulin molecule. The essential subunits of γ-TuRCs are 2-spoke γ-tubulin small complexes (γ-TuSCs), made from GCP2, GCP3 and two γ-tubulins. In budding yeast, γ-TuSCs are stimulated to assemble into helical structures when bound by the conserved “Centrosomin motif 1” (CM1) domain found within the yeast spindle pole body (SPB; centrosome equivalent) proteins Spd110 and Spc72 (Stu2, a TOG domain protein, is also required in the case of Spc72) (Kollman et al., 2010; Gunzelmann et al., 2018a). The “CM1 motif” within Spc110’s CM1 domain binds across adjacent γ-TuSCs, which presumably promotes the oligomerisation process at the SPB (Brilot et al., 2021; Lyon et al., 2016; Kollman et al., 2010; Lin et al., 2014). In most eukaryotes, however, γ-TuSCs are stimulated to assemble into γ-TuRCs within the cytosol via a 4-spoke GCP4/5/4/6 core complex that seeds ring assembly and is absent from budding yeast (Haren et al., 2020; Würtz et al., 2022). Indeed, depletion of GCP4, GCP5 or GCP6 strongly inhibits cytosolic γ-TuRC assembly in human, *Xenopus*, *Drosophila*, *Aspergillus*, and fission yeast cells (Cota et al., 2017; Vogt et al., 2006; Vérollet et al., 2006; Xiong and Oakley, 2009; Zhang et al., 2000). Intriguingly, however, these γ-TuRC-specific proteins are not essential in *Drosophila*, *Aspergillus*, or fission yeast (Xiong and Oakley, 2009; Vogt et al., 2006; Anders et al., 2006; Vérollet et al., 2006). Consistent with this, γ-TuSCs can be recruited to *Drosophila* S2 cell centrosomes after depletion of GCP4/5/4/6 core complex components (Vérollet et al., 2006), and are recruited independently of the GCP4/5/4/6 core complex to the outer SPB plaque in *Aspergillus* (Gao et al., 2019). Nevertheless, how γ-TuSCs are recruited to centrosomes in the absence of the GCP4/5/4/6 core remains unclear.

The predominant view of γ-TuRC recruitment involves the binding of large coiled-coil “tethering proteins” whose experimental depletion leads to measurable reductions in γ-tubulin levels at centrosomes. One of these proteins, NEDD1/Grip71, associates with pre-formed γ-TuRCs in the cytosol and subsequently docks γ-TuRCs to centrosomes (Lüders et al., 2006; Haren et al., 2006, 2009; Zhang et al., 2009; Gomez-Ferreria et al., 2012a; b). All other tethering proteins do not associate with cytosolic γ-TuRCs but instead localise to centrosomes and appear to “dock” incoming γ-TuRCs. These include the Pericentrin family of proteins, CM1 domain-containing proteins (e.g. human CDK5RAP2, *Drosophila* Centrosomin (Cnn), fission yeast Mto1, and budding yeast Spc110 and Spc72), and the Spd-2 family of proteins (CEP192 in humans) (Zimmerman et al., 2004; Gomez-Ferreria et al., 2007; Haren et al., 2009; Fong et al., 2008; Sawin et al., 2004; Zhang and Megraw, 2007; Conduit et al., 2014; Dobbelaere et al., 2008; Lee and Rhee, 2011). It is difficult to determine the individual role of these proteins in γ-TuRC recruitment, as they act redundantly and depend on each other for their proper localisation within the PCM. We previously showed that γ-tubulin partially accumulated at mitotic centrosomes in the absence of either Cnn or Spd-2, but failed to accumulate when both proteins were removed, with centrosomes also failing to accumulate other PCM proteins and nucleate microtubules (Conduit et al., 2014). This data showed that Cnn and Spd-2 can independently recruit γ-tubulin-containing complexes (hereafter γ-tubulin complexes), but it remained unclear how.

Cnn contains the highly conserved CM1 domain (Sawin et al., 2004), which binds directly to γ-tubulin complexes in yeast and humans, respectively, (Brilot et al., 2021; Wieczorek et al., 2019; Choi et al., 2010) and has been implicated in the recruitment of γ-tubulin complexes to centrosomes in different systems (Zhang and Megraw, 2007; Lyon et al., 2016; Samejima et al., 2008; Choi et al., 2010; Muroyama et al., 2016; Fong et al., 2008). But whether the CM1 domain is essential for Cnn to recruit γ-tubulin complexes remains unclear, as the effect of removing the CM1 domain has only been tested in the presence of Spd-2 (Zhang and Megraw, 2007). In contrast to Cnn, Spd-2 does not contain a CM1 domain and so how it recruits γ-tubulin complexes remains to be established.

In this study, we investigated how γ-tubulin complexes are recruited to mitotic centrosomes by Cnn and Spd-2. We used classical *Drosophila* genetics to combine specific mutant alleles or RNAi constructs and examined γ-tubulin accumulation at centrosomes in larval brain cells. We show that Cnn allows the centrosomal accumulation of γ-tubulin in the absence of the GPC4/5/4/6 core and Grip71 and that this is dependent on its CM1 domain. Mutations in the CM1 domain also abolish Cnn’s ability to associate with γ-tubulin in immunoprecipitation assays. This suggests that Cnn’s CM1 domain can bind and recruits γ-TuSCs to centrosomes, similar to the CM1 domains in budding yeast Spc110 and Spc72. In contrast, we find that Spd-2 does not support the centrosomal accumulation of γ-tubulin in the absence of the GPC4/5/4/6 core and Grip71, suggesting that Spd-2 can recruit only γ-TuRCs that have pre-formed in the cytosol. By selectively abolishing γ-tubulin complex recruitment, we show that mitotic centrosomes can nucleate microtubules independently of γ-tubulin complexes and that this depends on the TOG domain protein Mini-spindles (Msps), consistent with the conserved ability of TOG domain proteins to promote microtubule nucleation *in vitro*. Moreover, the microtubules nucleated in the absence of γ-TuRCs are more cold-stable than those nucleated in the presence of γ-TuRCs, suggesting that the dynamic properties of microtubules depend in part on how the microtubules were nucleated.

## Results

### Cnn recruits γ-tubulin complexes via its CM1 domain independently of Grip71 and the GCP4/5/4/6 core

We first explored how Cnn recruits γ-tubulin complexes to mitotic centrosomes by comparing the levels of centrosomal γ-tubulin at interphase and mitotic centrosomes in larval brain cells from flies depleted of Spd-2 and different γ-tubulin complex proteins. Typically, interphase centrosomes have only ∼5-20% of the γ-tubulin levels found at mitotic centrosomes, and this residual γ-tubulin is closely associated with the centrioles and is non-functional with respect to microtubule nucleation (Conduit et al., 2014). An increase in γ-tubulin signal between interphase and mitotic centrosomes indicates that γ-tubulin complexes have been recruited to the expanding mitotic PCM, at least to some degree. Similar to our previous results (Conduit et al., 2014), we found that γ-tubulin was recruited to mitotic centrosomes in *spd-2* null mutant brains with an average level of ∼77% compared to wild-type brains (Figure 1A,B). We know that this recruitment of γ-tubulin is entirely dependent on Cnn, because centrosomes in *cnn,spd2* double mutants fail entirely to recruit γ-tubulin during mitosis (Conduit et al., 2014). In contrast, combining *spd-2* null mutant alleles with null or severe depletion mutant alleles, or RNAi alleles, for Grip71 and the GCP4/5/4/6 core components did not prevent γ-tubulin accumulation at mitotic centrosomes. In fact, the centrosomes in *spd-2,grip71,grip75^GCP4^,grip128^GCP5^*-RNAi,*grip163^GCP6^* mutant cells had ∼66% of the γ-tubulin levels found at wild-type centrosomes, only slightly lower than ∼77% in *spd-2* mutants alone (Figure 1A,B). Thus, the recruitment of γ-tubulin to mitotic centrosomes that occurs in the absence of Spd-2, i.e. that depends upon Cnn, does not appear to require Grip71 or the GCP4/5/4/6 core.

**Figure 1.**
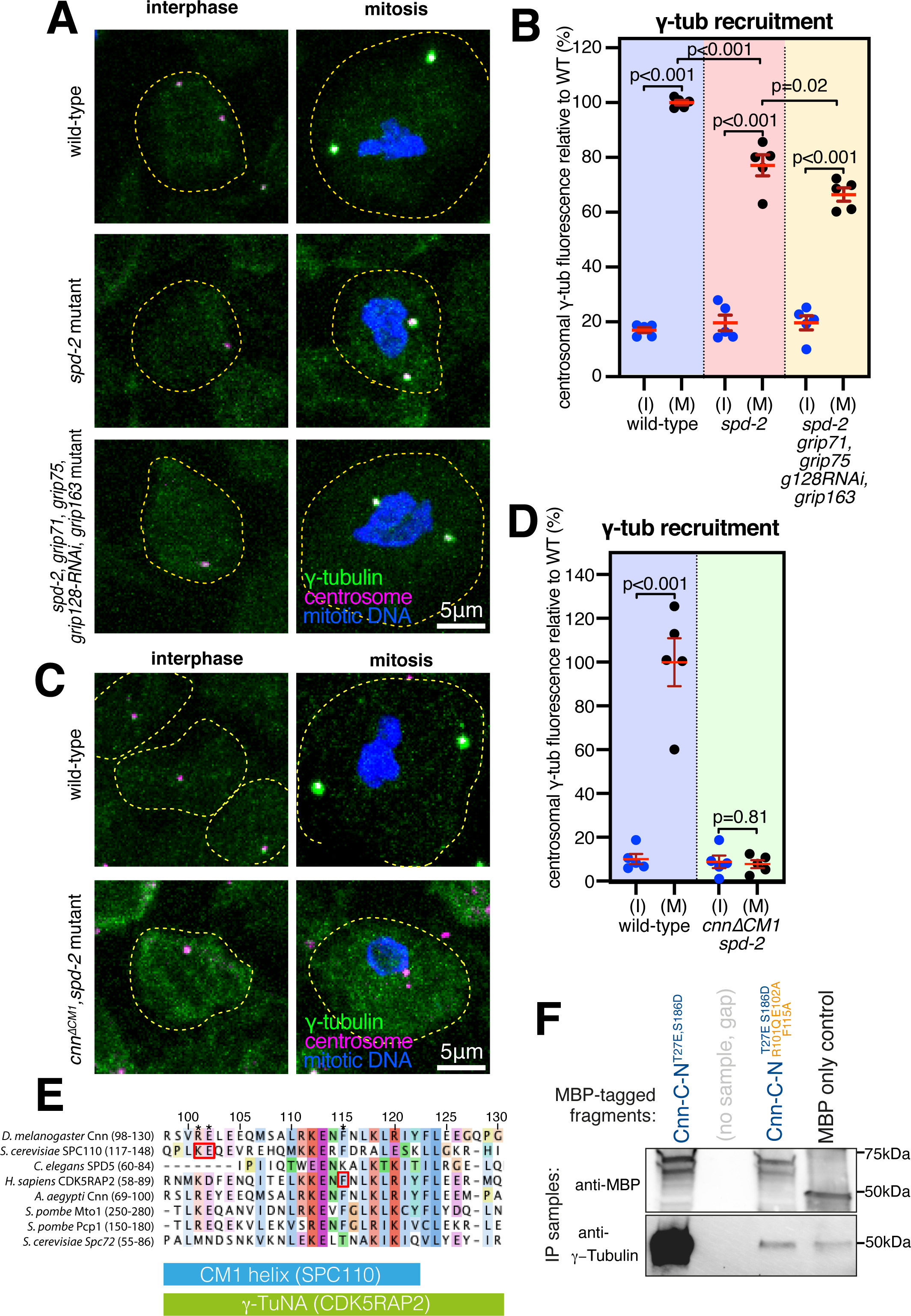
Cnn recruits γ-tubulin complexes via its CM1 domain and independently of Grip71 and the GCP4/5/4/6 core. **(A)** Fluorescence images of either interphase or mitotic *Drosophila* brain cells from either wild-type, *spd-2* mutant, or *spd-2, grip71,grip75^GCP4^,grip128^GCP5^*-RNAi,*grip163^GCP6^*mutant third instar larval brains immunostained for γ-tubulin (green), mitotic DNA (blue), and Asl (centrioles, magenta). Both mutants carry the mutant *spd-2* alleles, to reveal the Cnn pathway of recruitment. Scale bar is 5μm and applies to all images. **(B)** Graph showing average centrosomal fluorescence intensities of γ-tubulin (relative to wild-type) of interphase (blue dots) and mitotic (black dots) centrosomes from different genotypes (as indicated below). Each data-point represents the average centrosome value from one brain. Mean and SEM are indicated. A one-way ANOVA with a Sidak’s multiple comparisons test was used to make the comparisons indicated by p values in the graph. Note that there is only a small reduction in mitotic centrosomal γ-tubulin levels in *spd-2* mutants and in *spd-2, grip71,grip75^GCP4^, grip128^GCP5^*-RNAi,*grip163^GCP6^*mutants, showing that Cnn can still efficiently recruit γ-tubulin complexes to mitotic centrosomes when only γ-TuSCs are present. **(C)** Fluorescence images of either interphase or mitotic *Drosophila* brain cells from either wild-type or *cnn*^Δ*CM1*^,*spd-2* mutant third instar larval brains immunostained for γ-tubulin (green), mitotic DNA (blue), and Asl (centrioles, magenta). Scale bar is 5μm and applies to all images. **(D)** Graph in the same format as in (B) revealing no significant increase of centrosomal γ-tubulin signal from interphase to mitosis in *cnn*^Δ*CM1*^;*spd-2* mutant cells, showing that Cnn requires its CM1 domain to recruit γ-tubulin complexes to centrosomes. Ratio paired t-tests were used to compare mean values of interphase and mitotic centrosomes within each genotype. **(E)** Multi-protein sequence alignment of part of the CM1 domain containing the key binding residues (indicated by red boxes) in budding yeast and humans that we mutated in *Drosophila*. **(F)** Western blot probed for MBP and γ-tubulin showing the results of IP experiments from embryo extracts using bacterially purified MBP-tagged N-terminal (aa1-255) Cnn fragments containing point mutations to relieve Cnn-C autoinhibition (T27E and S186D; Tovey et al., 2021) and to perturb the CM1 domain’s ability to bind γ-TuRCs (R101Q, E102A, and F115A).

While we cannot rule out that residual amounts of GCP4/5/4/6 core components in *spd-2,grip71,grip75^GCP4^,grip128^GCP5^*-RNAi,*grip163^GCP6^*mutant cells may support a certain level of γ-TuSC oligomerisation in the cytosol, we favour the conclusion that Cnn can recruit γ-TuSCs directly to centrosomes in the absence of the GCP4/5/4/6 core for several reasons: First, the alleles used for *grip71* and *grip75^GCP4^*are null mutants, and the allele for *grip163^GCP6^* is a severe depletion allele (see Methods), and even individual mutations in, or RNAi-directed depletion of, Grip75^GCP4^, Grip128^GCP5^ or Grip163^GCP6^ are sufficient to strongly reduce the presence cytosolic γ-TuRCs (Vogt et al., 2006; Vérollet et al., 2006). Second, *spd-2,grip71,grip75^GCP4^,grip128^GCP5^*-RNAi,*grip163^GCP6^*mutant cells are depleted for all structural γ-TuRC components except for γ-TuSCs and Actin (note that Mozart1 (Mzt1) is not expressed in larval brain cells (Tovey et al., 2018) and that Mzt2 has not been identified in flies). In human and Xenopus γ-TuRCs, Actin supports γ-TuRC assembly via interactions with a GCP6-N-term-Mzt1 module (Liu et al., 2019; Wieczorek et al., 2019, 2020; Zimmermann et al., 2020; Consolati et al., 2020), and so Actin alone is unlikely to facilitate assembly of γ-TuSCs into higher order structures. Third, our data agree with the observation that near complete depletion of Grip71, Grip75^GCP4^, Grip128 ^GCP5^, and Grip163^GCP6^ from S2 cells does not prevent γ-tubulin recruitment to centrosomes (Vérollet et al., 2006). Fourth, given the strength of mutant alleles used, one would have expected a much larger decrease in centrosomal γ-tubulin levels in *spd-2,grip71,grip75^GCP4^,grip128^GCP5^*-RNAi,*grip163^GCP6^* mutant cells were Cnn not able to recruit γ-TuSCs directly to centrosomes. Thus, Cnn appears to recruit γ-TuSCs to centrosomes without a requirement for them to first assemble into higher-order complexes.

The recruitment of γ-TuSCs to centrosomes by Cnn appears to reflect the natural situation in budding yeast, where homologues of the GCP4/5/4/6 core, Grip71 and Mzt1 are absent and where γ-TuSCs are recruited to the SPB by direct binding of Spc110 and Spc72’s CM1 domain. We therefore reasoned that Cnn’s recruitment of γ-TuSCs may also be mediated by its CM1 domain. A previous study, however, had shown that replacing the endogenous *cnn* gene with an ectopically expressed UAS-GFP-Cnn construct lacking the CM1 domain led to a reduction, but not elimination, of γ-tubulin at centrosomes in syncytial embryos (Zhang and Megraw, 2007). Along with the potential effects caused by Cnn over-expression, we now know that Spd-2 can recruit γ-tubulin complexes independently of Cnn (Conduit et al., 2014), making it hard to evaluate the true effect of deleting the CM1 domain without also removing Spd-2. We therefore deleted the CM1 domain (amino acids 98-167, inclusive) from the endogenous *cnn* gene (see Methods) and combined this mutant allele with the *spd-2* null mutant allele. We found that γ-tubulin no longer accumulated at mitotic centrosomes in these *cnn*^Δ*CM1*^*,spd-2* mutant cells (Figure 1C,D), showing definitively that Cnn’s CM1 domain is essential for Cnn to recruit γ-tubulin complexes to mitotic centrosomes.

We also tested whether the CM1 domain was required for Cnn to associate with γ-tubulin complexes by comparing the ability of bacterially purified MBP-tagged Cnn fragments to co-immunoprecipitate γ-tubulin from wild-type cytosolic embryo extracts. We recently showed that Cnn’s centrosomal isoform (Cnn-C) is auto-inhibited from binding cytosolic γ-tubulin complexes by an extreme N-terminal “CM1 auto-inhibition” (CAI) domain, but that this auto-inhibition can be relieved by introducing T27E and S186D phospho-mimetic mutations (Tovey et al., 2021). These mutations were therefore included in the fragments to enable Cnn binding in “control” conditions (Cnn-C-N^T27E,S186D^). To identify mutations predicted to perturb CM1 binding, we used cross-species protein sequence alignments and identified F115, R101, and E102 as equivalent to residues important for γ-tubulin complex binding in humans (F75) (Choi et al., 2010) and budding yeast (K120 and E121) (Lin et al., 2014; Gunzelmann et al., 2018b) (Figure 1E). We mutated these residues in the Cnn-C-N^T27E,S186D^ fragments (R101Q, E102A and F115A) to mimic the mutations previously used in yeast and human experiments. Strikingly, these mutations abolished the ability of the Cnn fragments to co-immunoprecipitate γ-tubulin (Figure 1F), showing that Cnn’s CM1 domain is required for its binding to γ-tubulin complexes.

Taken together, our data strongly indicate that, similar to Spc110 and Spc72 in budding yeast, Cnn can bind and recruit γ-TuSCs to centrosomes directly from the cytosol without the need for them to pre-form into γ-TuRCs in the cytosol.

### Spd-2 predominantly recruits pre-formed γ-TuRCs from the cytosol

To explore how Spd-2 recruits γ-tubulin complexes even though (unlike Cnn) it lacks a CM1 domain, we compared the levels of centrosomal γ-tubulin at interphase and mitotic centrosomes in larval brain cells from flies lacking Cnn and different γ-tubulin complex proteins. In *cnn* mutants alone, Spd-2 levels at mitotic centrosomes are reduced to ∼60% (Conduit et al., 2014) and this Cnn-independent pool of Spd-2 recruits γ-tubulin to on average ∼22-23% of wild-type levels (Figure 2A,B) (Conduit et al., 2014). We were therefore testing which γ-TuRC components, when removed in addition to Cnn, reduced the mitotic centrosomal level of γ-tubulin further, such that there was no significant accumulation of γ-tubulin above interphase levels. Of note, the Cnn-dependent pool of Spd-2 also recruits some γ-tubulin complexes, because γ-tubulin levels were slightly higher at centrosomes in Cnn^ΔCM1^ mutant cells compared to *cnn* null mutant cells (Figure S1A,B). The absence of a large increase may be because deleting the CM1 domain could affect the ability of Cnn to form a robust scaffold and therefore maintain Spd-2 in the PCM, as we noticed that the γ-tubulin signals were often offset from the centriole signal (Figure S1A), as it is in cnn mutant cells (Figure 2A; (Figure S1A) (Lucas and Raff, 2007)).

**Figure 2.**
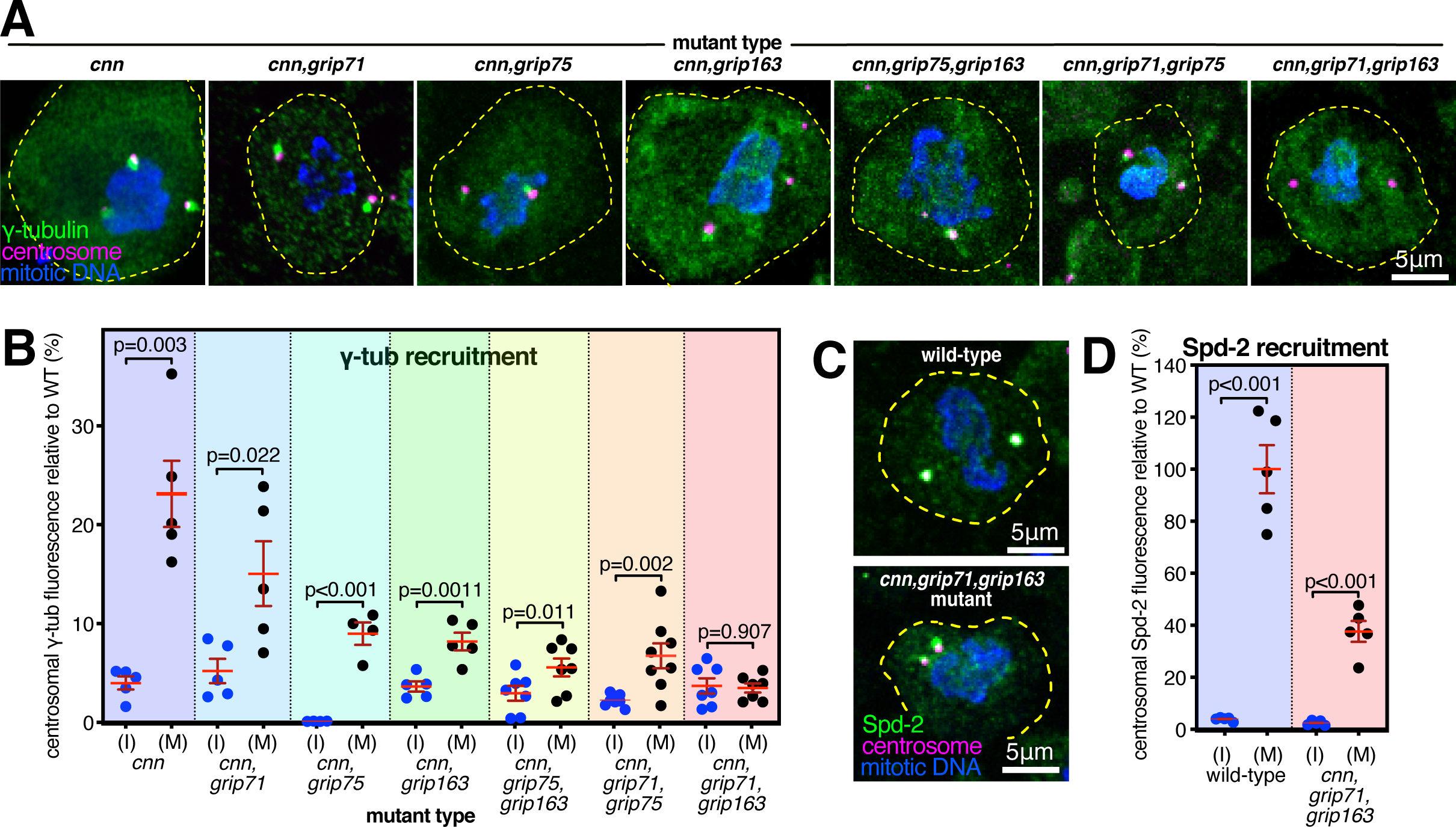
Recruitment of γ-tubulin complexes by Spd-2 is heavily dependent on the GCP4/5/4/6 core. **(A)** Fluorescence images of mitotic *Drosophila* brain cells from various mutant third instar larvae immunostained for γ-tubulin (green), mitotic DNA (blue), and Asl (centrioles, magenta). All mutants carry the mutant *cnn* allele to reveal the Spd-2 pathway of recruitment, along with mutant alleles for different combinations of γ-TuRC genes. Scale bar is 5μm and applies to all images. **(B)** Graph showing average centrosomal fluorescence intensities of γ-tubulin (relative to wild-type) of interphase (blue dots) and mitotic (black dots) centrosomes from different genotypes (as indicated below). Each data-point represents the average centrosome value from one brain. Mean and SEM are indicated. Paired t-tests were used to compare mean values of interphase and mitotic centrosomes within each genotype. Note that γ-tubulin accumulation at mitotic centrosomes is severely perturbed in the absence of the GCP4/5/4/6 core components Grip75^GCP4^ and Grip163^GCP6^, but is abolished only in the absence of Grip71 and Grip163^GCP6^. **(C)** Fluorescence images of mitotic *Drosophila* brain cells from wild-type or *cnn,grip71,grip163* mutant third instar larvae immunostained for Spd-2 (green), mitotic DNA (blue), and Asl (centrioles, magenta). Scale bar is 5μm and applies to both images. **(D)** Graph in the same format as in (B) showing a significant increase of centrosomal Spd-2 signal from interphase to mitosis in *cnn,grip71,grip163* mutant cells, showing that the inability of these centrosomes to recruit γ-tubulin (A,B) is not due to an absence of Spd-2.

We predicted that Spd-2 recruits γ-tubulin complexes via Grip71 because the human homologue of Spd-2, Cep192, associates with the human homologue of Grip71, NEDD1 (Gomez-Ferreria et al., 2012a). We found, however, that γ-tubulin could still accumulate at mitotic centrosomes relatively well in *cnn,grip71* mutant cells, being only slightly lower than the γ-tubulin levels in *cnn* mutant cells (Figure 2A,B). Thus, Spd-2 relies only partly on Grip71 to recruit γ-tubulin complexes. There was a stronger reduction, however, when we removed Cnn and members of the GCP4/5/4/6 core, Grip75^GCP4^ and Grip163^GCP6^, either individually or in combination (Figure 2A,B). Given that the GCP4/5/4/6 core is required for the assembly of γ-TuRCs within the cytosol, this result suggests that Spd-2 (unlike Cnn) predominantly recruits pre-formed γ-TuRCs rather than γ-TuSCs. We found, however, that γ-tubulin accumulation at mitotic centrosomes was only abolished after the additional removal of Grip71 i.e. in *cnn,grip71,grip163^GCP6^* mutant cells (Figure 2A,B), a phenotype that was not due to a failure of Spd-2 to accumulate at mitotic centrosomes (Figure 2C,D). Thus, Spd-2 appears to recruit a very small amount of γ-TuSCs (which may, or may not, be present as larger assemblies due to an association with Grip128-γ-tubulin) via Grip71 (i.e. the recruitment that occurs in cnn,grip75^GCP4^,grip163 ^GCP6^ cells), but its recruitment of γ-tubulin complexes relies predominantly on the GCP4/5/4/6 core.

Intriguingly, γ-tubulin could still accumulate at mitotic centrosomes to some degree in cells lacking Cnn, Grip71 and Grip75^GCP4^, showing that removal of Grip75^GCP4^ does not perfectly phenocopy the removal of Grip163^GCP6^, and therefore suggesting that Grip163^GCP6^ may still be able to promote at least partial γ-TuRC assembly in the absence of Grip75^GCP4^. This is consistent with observations in human cells, where GCP6 depletion has a greater effect on cytosolic γ-TuRC assembly than GCP4 depletion (Cota et al., 2017). Alternatively, Spd-2 may interact with Grip163^GCP6^ and so be able to recruit its associated γ-tubulin independent of Grip75^GCP4^.

In summary, Spd-2’s recruitment of γ-TuRCs relies strongly on the presence of the GCP4/5/4/6 core, and therefore on γ-TuRC assembly within the cytosol, but the additional removal of Grip71 is required to entirely prevent accumulation of γ-tubulin at mitotic centrosomes. In contrast, Cnn’s conserved CM1 domain can mediate the recruitment of γ-TuSCs directly from the cytosol. The requirement of Spd-2 for the GCP4/5/4/6 core aligns with the absence of Spd-2 and GCP4/5/4/6 core component homologues in lower eukaryotes. In addition, the ability of Cnn to recruit γ-TuSCs may explain why the GCP4/5/4/6 core is not essential in several species studied so far, particularly if all CM1 domain proteins are able to stimulate γ-TuSC assembly into ring-like structures, as is the case for yeast Spc110 and Spc72.

### Centrosomes lacking γ-tubulin can still nucleate and organise microtubules

In the course of examining *cnn,grip71,grip163* mutants, we observed that their mitotic centrosomes, which fail to accumulate γ-tubulin but still accumulate Spd-2, were still associated with microtubules during prophase and localised to spindle poles during mitosis (Figure 3A). This is in contrast to centrosomes in *cnn,spd-2* mutant cells, which lack PCM entirely, fail to nucleate or organise microtubules, and do not associate with spindle poles (Conduit et al., 2014). Thus, mitotic centrosomes can organise microtubules independently of γ-TuRCs so long as the PCM can at least partially assemble. To test whether these microtubules are actually nucleated at centrosomes (rather than being nucleated elsewhere and then attaching to the centrosomes) we performed a cooling-warming microtubule repolymerisation assay. We depolymerised microtubules by cooling larval brains on ice for ∼40 minutes and then either chemically fixed samples on ice or allowed them to warm up for 30 seconds before rapid chemical fixation. ∼40 minutes of cooling efficiently depolymerised microtubules at most centrosomes in wild-type and *cnn,grip71,grip163* mutant centrosomes (Figure 3B). After 30s warming, all wild-type and *cnn,grip71,grip163* centrosomes had an associated α-tubulin signal, either as asters or as part of a re-formed mitotic spindle (Figure 3C), strongly suggesting that an accumulation of γ-tubulin at mitotic centrosomes is not necessary for these centrosomes to nucleate microtubules.

**Figure 3.**
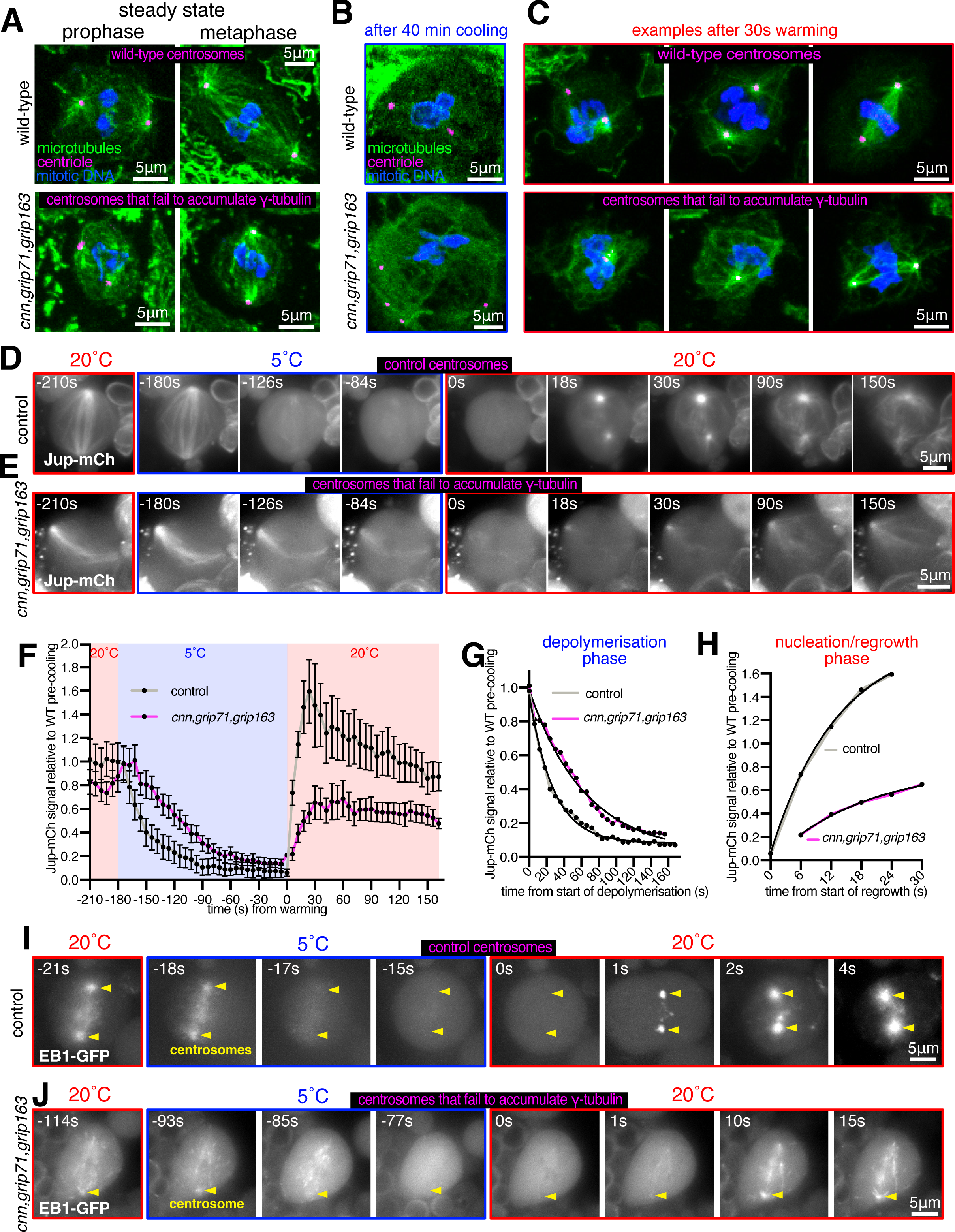
Mitotic centrosomes that fail to accumulate γ-tubulin can still nucleate microtubules. **(A-C)** Fluorescence images of mitotic *Drosophila* brain cells from either wild-type or *cnn,grip71,grip163* mutant third instar larval brains, either at steady state (A), after 40 minutes of cooling on ice (B), or after 30s of warming (post cooling) to room temperature (C) immunostained for alpha-tubulin (microtubules, green), mitotic DNA (blue), and Asl (centrioles, magenta). Note how centrosomes in both wild-type and *cnn,grip71,grip163* mutant cells are associated with microtubules both at steady state and after 30s warming. Note that some cells lacking Cnn have abnormal numbers of centrosomes due to centrosome segregation problems during cell division (Conduit et al., 2010). (**D-F**) Fluorescent images (D,E) and graph (F) documenting the behaviour of the microtubule marker Jupiter-mCherry within living *Drosophila* control (D) or *cnn,grip71,grip163* mutant (E) third instar larval brain cells as they were cooled to 5°C for ∼3 minutes and then rapidly warmed to 20°C. Time in seconds relative to the initiation of warming (0s) is indicated. Note that the GFP-PACT signal used to locate centrosomes is not displayed. The graph in (F) plots the mean and SEM centrosomal signal (after subtraction of cytosolic signal) of 12 and 10 centrosomes from 7 and 10 control and *cnn,grip71,grip163* mutant cells, respectively. The data is normalised to the average signal at centrosomes in control cells prior to cooling. Note how the centrosomal Jupiter-mCherry signal quickly drops on cooling and then immediately increases on warming in both control and *cnn,grip71,grip163* mutant cells, showing that centrosomes within both control and *cnn,grip71,grip163* mutant cells nucleate microtubules. (**G,H**) Graphs show the depolymerisation (G) and nucleation/regrowth phases (H) phases from the graph in (F). One-phase exponential decay models and “exponential plateau” models generated in GraphPad Prism are fitted to the depolymerisation and nucleation/regrowth phases, respectively. Note how the centrosomal Jupiter-mCherry signal decreases faster upon cooling, but increases slower upon warming, in *cnn,grip71,grip163* mutant cells. (**I,J**) Fluorescent images documenting the behaviour of the microtubule plus-end marker EB1-GFP within living *Drosophila* control (I) and *cnn,grip71,grip163* mutant (J) third instar larval brain cells as they were cooled to 5°C and then rapidly warmed to 20°C. Time in seconds relative to the initiation of warming (0s) is indicated. Note how the EB1-GFP signal emanates from the centrosome and from the spindle/chromatin region during warming in the *cnn,grip71,grip163* mutant cell.

To better understand microtubule dynamics at wild-type and *cnn,grip71,grip163* centrosomes we established a system to image cells live while cooling and warming the sample. We generated stocks containing fluorescent markers of microtubules (Jupiter-mCherry) and centrosomes (GFP-PACT) with and without the *cnn*, *grip71* and *grip163* mutations and used a microscope-fitted heating-cooling device (CherryTemp) to modulate the temperature of larval brain samples during recording. We imaged samples for ∼30s before cooling them to 5°C for 3 minutes to depolymerise microtubules and then rapidly warming them to 20°C to observe microtubule regrowth. When cells were cooled to 5°C, the centrosomal Jupiter-mCherry signal decreased towards cytosolic background levels at both control and *cnn,grip71,grip163* centrosomes (Figure 3D-F; Videos S1 and S2). In a subset of cells, this centrosomal signal reached cytosolic levels (i.e. disappeared) after 3 minutes of cooling (Figure S3A,B). On warming to 20°C, there was an immediate increase in the centrosomal Jupiter-mCherry signal at all control and *cnn,grip71,grip163* centrosomes (Figure 3D-F; Figure S3A,B), confirming that microtubules can be nucleated at mitotic centrosomes that have not accumulated γ-tubulin. The dynamics of the microtubules differed, however (Figure 3F). On cooling to 5°C, microtubules depolymerised faster at control centrosomes (Figure 3F) – fitting “one-phase exponential decay” models to the depolymerisation phases produced half-lives of 21.79 and 44.88 seconds and decay rate constants of 0.0361 and 0.0154 for control and *cnn,grip71,grip163* centrosomes, respectively. On warming to 20°C, microtubules also polymerised faster at control centrosomes (Figure 3F) – fitting “exponential plateau” models produced growth rate constants of 0.0759 and 0.0536, respectively, which when normalised to the YM values (the maximum plateau values) showed an ∼3.4-fold difference in growth rate (Figure 3H). Differences in microtubule dynamics were also apparent when imaging EB1-GFP comets, which mark growing microtubule plus ends. EB1-GFP comets emerging from control centrosomes disappeared immediately after cooling to 5°C and then reappeared immediately after warming to 20°C (Figure 3I; Video S3), but comets emerging from mutant centrosomes took longer to disappear, and fewer reappeared, during cooling and warming cycles (Figure 3J; Video S4). Moreover, it was easier to observe EB1-GFP comets emerging from chromatin regions in these *cnn,grip71;grip163* mutant cells (Figure 3J; Video S4), presumably because the centrosomes were no longer such dominant sites of microtubule nucleation. Thus, microtubules depolymerise faster and are then nucleated and/or polymerised faster at control centrosomes compared to at *cnn,grip71,grip163* centrosomes.

One caveat with the experiments above is that centrosome assembly is strongly perturbed in cells lacking the centrosome scaffold protein Cnn (Lucas and Raff, 2007; Conduit et al., 2014), potentially impacting the γ-TuRC-independent ability of centrosomes in *cnn,grip71;grip163* mutant cells to nucleate and organise microtubules. We therefore generated *cnn*^Δ*CM1*^*, grip71, grip163* mutants with or without GFP-PACT and Jupiter-mCherry, allowing us to examine microtubule dynamics at mitotic centrosomes that did not accumulate γ-tubulin (Figure 4A,B) but that still had Cnn to help assemble the PCM (although we note that PCM assembly appears perturbed to some degree in Cnn^ΔCM1^ mutant cells – see Figure S1). Prior to cooling, centrosomes in *cnn*^Δ*CM1*^*,grip71,grip163* mutant cells had a Jupiter-mCherry signal that was on average slightly higher than in control cells, suggesting robust microtubule organisation (Figure 4C-E). Similar to *cnn,grip71,grip163* mutants, microtubules depolymerised slower at *cnn*^Δ*CM1*^*,grip71,grip163* centrosomes compared to controls (Figure 4C-E). Fitting models to the data revealed half-lives of 16.7 and 37.7, and decay rate constants 0.0416 of 0.0184 for control and *cnn*^Δ*CM1*^*, grip71, grip163* mutants, respectively (Figure 4G). However, the signal plateaued at a relatively high value despite cooling for 5 minutes as opposed to 3 (Figure 4E), suggesting that a larger proportion of microtubules are cold-stable at *cnn*^Δ*CM1*^*,grip71,grip163* mutant centrosomes compared to *cnn*^Δ*CM1*^*,grip71,grip163* mutant and control centrosomes. On warming to 20°C, microtubules polymerised at *cnn*^Δ*CM1*^*,grip71,grip163* centrosomes but again at a slower rate than at control centrosomes (Figure 4C-E): growth rate constants normalised to the YM values showed an ∼3.4-fold difference in growth rate (Figure 4H), very similar to the growth rate for *cnn,grip71,grip163* mutant centrosomes.

**Figure 4.**
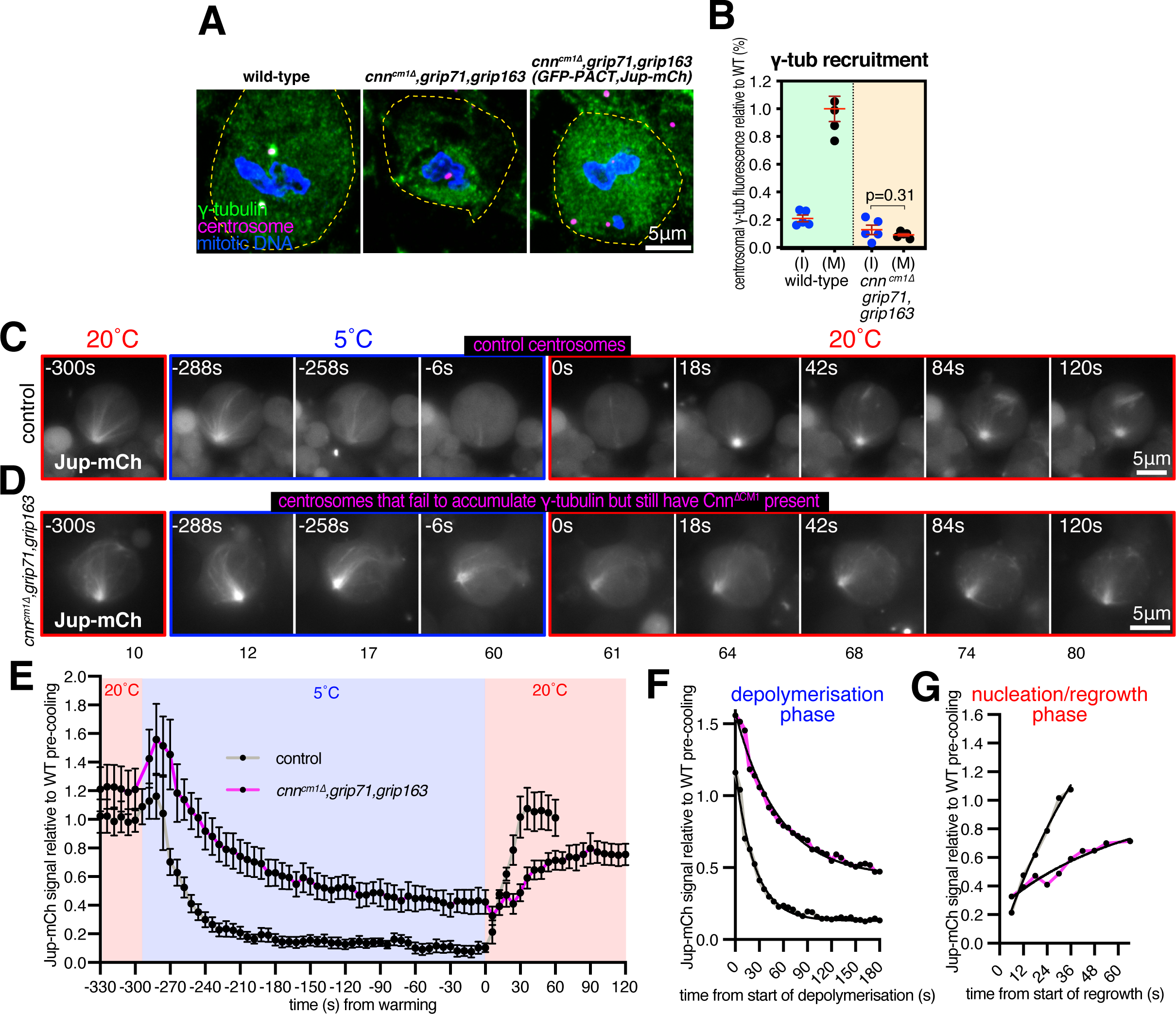
Mitotic centrosomes that fail to accumulate γ-tubulin nucleate microtubules that are cold-resistant. **(A-C)** Fluorescence images of mitotic *Drosophila* brain cells from either wild-type third instar larvae, *cnn*^Δ*CM1*^*,grip71,grip163* mutant third instar larvae, or *cnn*^Δ*CM1*^*,grip71,grip163* mutant third instar larvae also expressing GFP-PACT and Jupiter-mCherry, immunostained for γ-tubulin (green), mitotic DNA (blue), and Asl (centrioles, magenta). Note that GFP and mCherry fluorescence signals are destroyed during the fixation process due to addition of acetic acid. Scale bar is 5μm and applies to all images. **(B)** Graph showing average centrosomal fluorescence intensities of γ-tubulin (relative to wild-type) of interphase (blue dots) and mitotic (black dots) centrosomes from either wild-type or *cnn*^Δ*CM1*^*,grip71,grip163* mutants. Each data-point represents the average centrosome value from one brain. Mean and SEM are indicated. A Paired t-test was used to compare mean values of interphase and mitotic centrosomes, showing that there is no accumulation of γ-tubulin at mitotic centrosomes within the *cnn*^Δ*CM1*^*,grip71,grip163* mutant genotype. (**C-E**) Fluorescent images (C,D) and graph (E) documenting the behaviour of Jupiter-mCherry within living *Drosophila* control (C) or *cnn*^Δ*CM1*^*,grip71,grip163* mutant (D) third instar larval brain cells as they were cooled to 5°C for 5 minutes and then rapidly warmed to 20°C. Time in seconds relative to the initiation of warming (0s) is indicated. Note that the GFP-PACT signal used to locate centrosomes is not displayed. The graph in (E) plots the mean and SEM centrosomal signal (after subtraction of cytosolic signal) of 12 and 11 centrosomes from 4 and 4 control and *cnn*^Δ*CM1*^*,grip71,grip163* mutant cells, respectively. The data is normalised to the average signal at centrosomes in control cells prior to cooling. Note that a relatively large fraction of the centrosomal Jupiter-mCherry signal remains at centrosomes during cooling in *cnn*^Δ*CM1*^*,grip71,grip163* mutant cells, showing that the microtubule nucleated by these centrosomes are very cold-resistant. (**F,G**) Graphs show the depolymerisation (F) and nucleation/regrowth phases (G) phases from the graph in (E). One-phase exponential decay models and “exponential plateau” models generated in GraphPad Prism are fitted. Note how the centrosomal Jupiter-mCherry signal decreases faster upon cooling, but increases slower upon warming, in *cnn*^Δ*CM1*^*,grip71,grip163* mutant cells.

We conclude that centrosomes can nucleate microtubules independently of γ-TuRCs but that these microtubules are nucleated slower or grow slower and are more cold-stable than microtubules nucleated from wild-type centrosomes. This suggests that different modes of microtubule nucleation generate microtubules with different properties.

### The TOG domain protein Msps promotes microtubule nucleation from centrosomes lacking γ-tubulin complexes

We next addressed which proteins promote γ-TuRC-independent microtubule nucleation at mitotic centrosomes. We did not observe any clear enrichment of α-tubulin at centrosomes after microtubule depolymerisation (Figure 3B; Figure S2), ruling out the possibility that a high local concentration of α/β-tubulin accounts for or contributes to γ-TuRC-independent microtubule nucleation. Proteins of the chTOG/XMAP215 and TPX2 protein families have been reported to promote γ-TuRC-independent microtubule nucleation. These proteins promote microtubule nucleation in a range of species both *in vitro* and *in vivo*, including in the absence of γ-TuRCs (see Discussion and references therein). The *Drosophila* homologue of chTOG is Minispindles (Msps), which binds microtubules, localises to centrosomes and spindle microtubules and is required for proper spindle formation, mitotic progression and chromosome segregation (Cullen et al., 1999). Msps has also been reported to stabilise the minus ends of microtubules when bound and recruited to centrosomes by TACC (Barros et al., 2005; Lee et al., 2001). Msps is also part of a group of proteins that organise microtubules independently of γ-tubulin at the nuclear envelope of fat body cells (Zheng et al., 2020). Moreover, the TOG1 and 2 domains of Msps promote microtubule nucleation *in vitro* (Slep and Vale, 2007). The putative *Drosophila* TPX2 homologue is Mei-38 and, while its depletion results in only mild spindle defects, Mei-38 binds microtubules, localises to centrosomes and spindle microtubules, and is required for microtubule re-growth from kinetochores (Popova et al., 2022; Goshima, 2011). CAMSAP/Patronin/Nezha protein family members have also been implicated in γ-TuRC-independent microtubule nucleation and organisation at non-centrosomal sites (Akhmanova and Kapitein, 2022) and CAMSAP2 condensates can stimulate microtubule nucleation *in vitro* (Imasaki et al., 2022).

To test the role of these proteins in γ-TuRC-independent nucleation from centrosomes, we combined mutant or RNAi alleles with the *cnn*, *grip71*, and *grip163* mutant alleles and analysed microtubule organisation at centrosomes during prophase, when microtubule asters are most robust (Conduit et al., 2014). We were unable to obtain 3^rd^ instar lavae when combining the *cnn*, *grip71*, and *grip163* mutant alleles with *patronin* mutant or RNAi alleles, presumably due to severe microtubule defects that prevented development, and thus could not test the role of Patronin. We could, however, obtain larvae when combining the *cnn*, *grip71*, and *grip163* mutant alleles with mutant alleles for *msps* or *tacc* or an RNAi allele for Mei38. Clear association of microtubules with centrosomes was observed in 100% of wild-type prophase cells and in 96.8% of *cnn,grip71,grip163* mutant prophase cells (Figure 5A,B,F), consistent with our observations above that *cnn,grip71,grip163* centrosomes can nucleate and organise microtubules. In contrast, a clear association of microtubules with centrosomes was observed in only 55.3% of *cnn,grip71,grip163,msps* mutant prophase cells, and in 70.7% and 81.4% of *cnn,grip71,grip163,tacc* and *cnn,grip71,grip163*,*mei-38-RNAi* mutant cells, respectively (Figure 5C-F). Moreover, the *cnn,grip71,grip163,msps* centrosomes tended to be positioned further from the spindle poles than the *cnn,grip71,grip163* centrosomes (Figure 5G,H), which is indicative of a reduced capacity to organise microtubules. Positioning of centrosomes in *cnn,grip71,grip163,tacc* and *cnn,grip71,grip163*,*mei-38-RNAi* mutant cells was less affected, presumably due to the less severe defects in microtubule organisation at *cnn,grip71,grip163,tacc* and *cnn,grip71,grip163,Mei38-RNAi* centrosomes.

**Figure 5.**
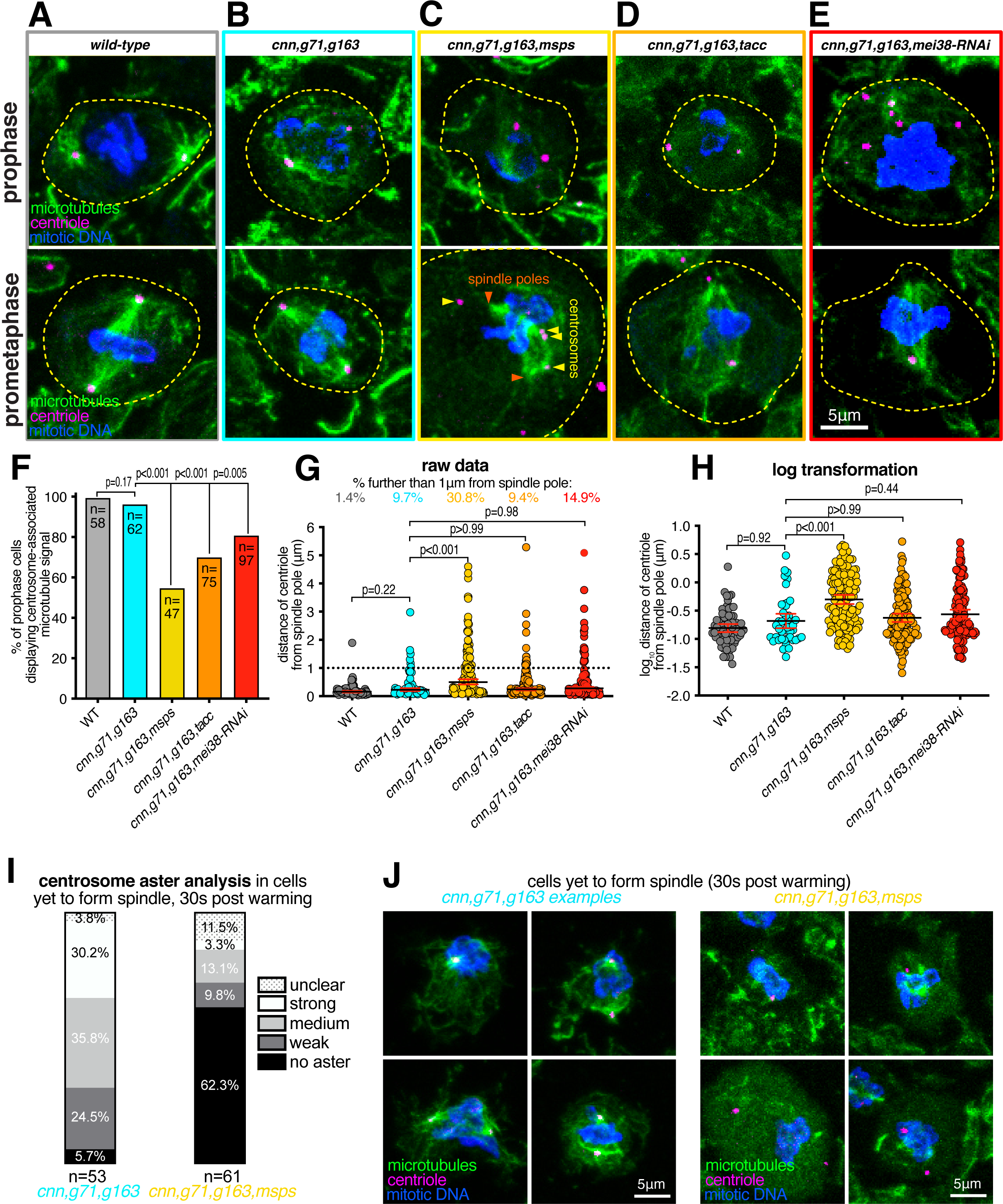
Depletion of Msps strongly perturbs the ability of centrosomes to organise and nucleate microtubules in the absence of γ-tubulin complexes. **(A-E)** Fluorescence images of *Drosophila* brain cells in either prophase or prometaphase from either wild-type (A), *cnn,grip71,grip163* (B), *cnn,grip71,grip163,msps* (C), *cnn,grip71,grip163,tacc* (D), or *cnn,grip71,grip163,mei38-RNAi* cell (E) immunostained for alpha-tubulin (microtubules, green), mitotic DNA (blue), and Asl (centrioles, magenta). Note that some cells lacking Cnn have abnormal numbers of centrosomes due to centrosome segregation problems during cell division (Conduit et al., 2010). **(F)** Graph showing the percentage of prophase cells in which microtubules are associated with at least one centrosome within the various genotypes, as indicated. The number of cells analysed (n) is indicated. Datasets were compared to the *cnn,grip71,grip163* dataset using one-way Chi-squared tests. Note how there is a no significant reduction between wild-type and *cnn,grip71,grip163* mutant cells in the proportion of cells displaying centrosome-associated microtubules, but there are significant reductions between *cnn,grip71,grip163* mutant cells and cells that are also depleted for either Msps, TACC or *mei*-38, indicating that Msps, TACC and *mei*-38 have a role in γ-TuRC-independent microtubule nucleation. **(G, H)** Graphs of raw data (G) and log transformed data (H) showing the distance of centrosomes from spindle poles during prometaphase in the different genotypes, as indicated. The percentage of centrosomes that were further than 1μm from the spindle poles is indicated above each dataset in the graph in (G). Increased distance from the spindle pole is indicative of a failure to properly organise microtubules. Kruskal-Wallis tests were used to compare the distribution of the *cnn,grip71,grip163* dataset with those of the other genotypes. Note that there is a significant difference only between *cnn,grip71,grip163* and *cnn,grip71,grip163,msps* mutant cells, indicating that Msps is particularly important for microtubule organisation at centrosomes in the absence of γ-TuRCs. **(I,J)** Parts of a whole graph (I) and images (J) represent analyses of centrosomal aster types in cells fixed and immunostained for alpha-tubulin (microtubules, green), mitotic DNA (blue), and Asl (centrioles, magenta) after 30 seconds of warming post cooling from either *cnn,grip71,grip163* or *cnn,grip71,grip163,msps* mutants, as indicated. Only cells that had centrosomes but that had not yet formed a spindle were analysed. Note how centrosome asters are frequently absent in *cnn,grip71,grip163,msps*.

Given that Msps appeared to be most important for γ-TuRC-independent nucleation of microtubules from centrosomes, we tested its role directly by performing a cooling/warming microtubule nucleation assay (similar to the fixed cell assay performed in Figure 3B,C) and compared the recovery of microtubules 30 seconds post warming at *cnn,grip71,grip163* centrosomes and at *cnn,grip71,grip163,msps* centrosomes. We categorised cells as those with or without centrosomes (some cells lack centrosomes due to mis-segregation of centrosomes during mitosis) and those that had or had not yet formed spindles; the proportion of these categories was similar in both mutant types (Figure S4). There were, however, differences between the mutant types within each category. Of the cells that contained centrosomes but had not yet formed a spindle, centrosomes organised microtubules in ∼94.3% of *cnn,grip71,grip163* mutant cells, the majority of which were scored as having strong or medium asters, but centrosomes organised microtubules in only ∼37.7% of *cnn,grip71,grip163,msps* mutant cells, the majority of which were scored as having weak asters (Figure 5I,J). This difference appeared to affect spindle formation because, of the cells that had centrosomes and that had formed a spindle structure, spindles were scored as being of “high” or “medium” quality (based on their morphology and density) in ∼67.1% of *cnn,grip71,grip163* mutant cells but in only ∼28% of *cnn,grip71,grip163,msps* mutant cells (Figure S5A,B). This was specific to centrosomes because there was a similarly high proportion of cells containing low quality spindles in both mutant types when cells lacked centrosomes (Figure S5C). For comparison, spindles were scored as being of “high” or “medium” quality in ∼95.3% of wild-type cells (Figure S5A). Note also that the absence of Grip71 abolishes the Augmin-mediated nucleation pathway necessary for efficient spindle assembly (Reschen et al., 2012; Chen et al., 2017b; Dobbelaere et al., 2008; Vérollet et al., 2006), but as both mutant types lacked Grip71 this cannot explain the differences observed between the mutants.

In summary, our data show that centrosomes lacking γ-tubulin complexes can still nucleate microtubules, despite having reduced PCM, and that the TOG domain protein Msps plays an important role in this γ-TuRC-independent microtubule nucleation pathway.

## Discussion

How centrosomes nucleate and organise microtubules is a long-standing question. Centrosomes contain hundreds of proteins, many of which associate with microtubules, meaning that understanding how centrosomes nucleate and organise microtubules is not trivial. Prior to our current work, we had identified Cnn and Spd-2 as the two key PCM components in flies – remove one and the other could support partial PCM assembly and microtubule organisation; remove both and PCM assembly and microtubule organisation fail (Conduit et al., 2014). We had found that γ-tubulin could still accumulate at mitotic centrosomes after removal of either Cnn or Spd-2, showing that both proteins could mediate the recruitment of γ-tubulin complexes, but it remained unclear how. The work presented here shows that Cnn and Spd-2 recruit different types of γ-tubulin complex, with Cnn able to recruit γ-TuSCs and Spd-2 recruiting predominantly pre-formed γ-TuRCs. Moreover, by preventing γ-tubulin recruitment but not PCM assembly, we have shown that centrosomes still nucleate microtubules in the absence of γ-TuRCs and that this γ-TuRC-independent mode of microtubule nucleation is stimulated by the TOG domain protein Msps (Figure 6).

**Figure 6.**
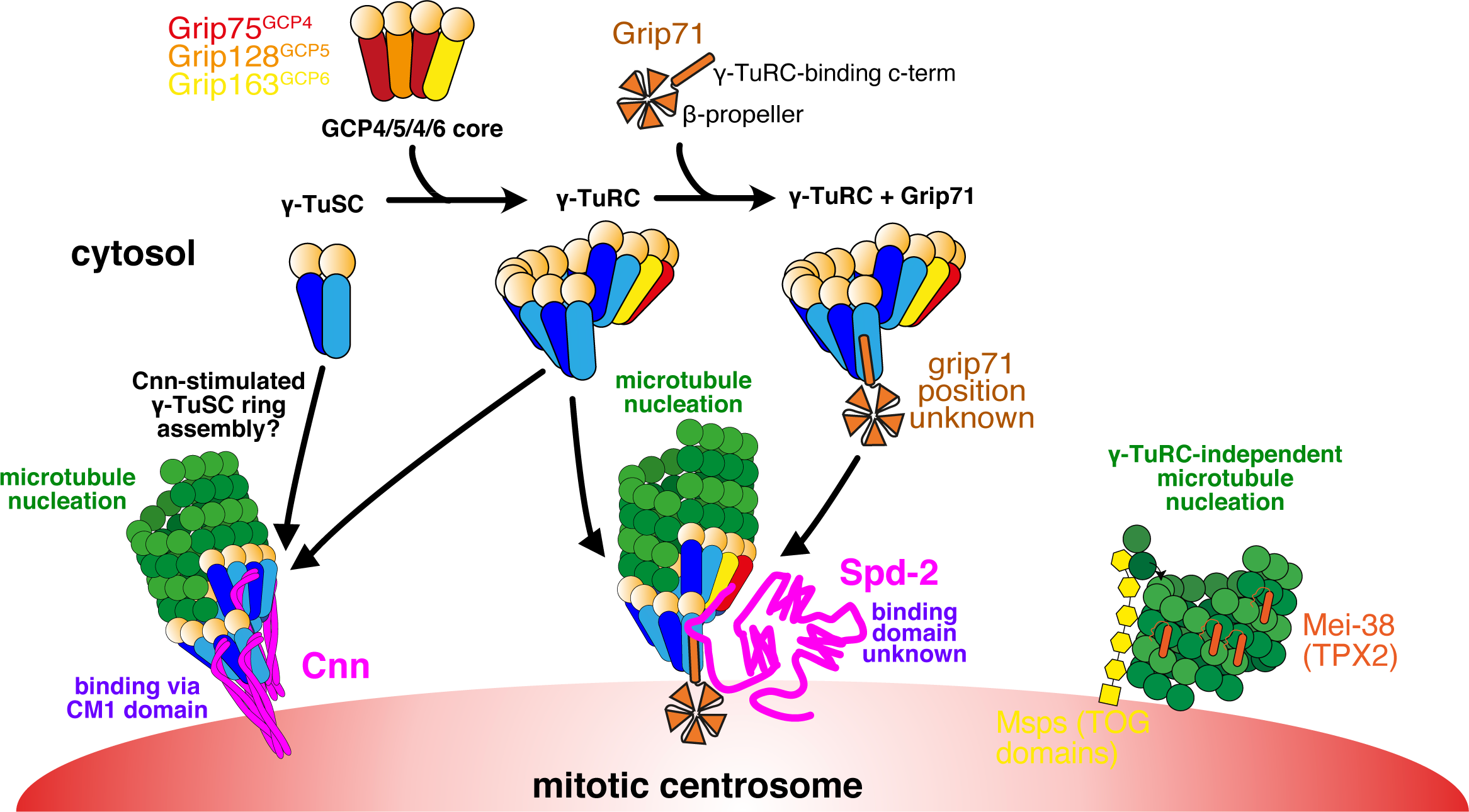
Model for the different pathways of γ-tubulin complex recruitment and microtubule nucleation at mitotic centrosomes in *Drosophila*. This model is based on both previous data from the literature and our current findings. A mixture of γ-TuSCs and γ-TuRCs exist in the cytosol. The GCP4/5/4/6 core, predicted to comprise Grip75^GCP4^, Grip128^GCP5^, and Grip163^GCP6^ in *Drosophila*, is necessary for γ-TuSCs to assemble into γ-TuRCs within the cytosol. Grip71 is a peripheral γ-TuRC protein that can associate with cytosolic γ-TuRCs but is not necessary for their assembly and so cytosolic γ-TuRCs with and without Grip71 may exist. Cnn is able to recruit γ-tubulin in the absence, or at least near absence, of the GCP4/5/4/6 core and Grip71, suggesting it can recruit γ-TuSCs directly from the cytosol. It likely also recruits pre-formed γ-TuRCs under normal conditions because artificial Cnn scaffolds recruit Grip75^GCP4^-GFP (Tovey et al., 2021). Cnn’s ability to recruit γ-tubulin complexes relies on its highly conserved N-terminal CM1 domain. We speculate that CM1 domain binding may stimulate γ-TuSC oligomerisation into γ-TuSC-only γ-TuRCs that could then nucleate microtubules, as is true of CM1 domain proteins in yeast. In contrast to Cnn, Spd-2 recruitment relies largely on the GCP4/5/4/6 core and so Spd-2 must predominantly recruit pre-formed γ-TuRCs. Spd-2 may be able to recruit very low levels of γ-TuSCs via Grip71 (not depicted). How Spd-2 binds to γ-tubulin complexes remains unknown. When the recruitment of γ-tubulin complexes by both Cnn and Spd-2 is inhibited, centrosomes are still able to nucleate microtubules and this γ-tubulin-independent microtubule nucleation pathway depends on Msps (the fly TOG domain protein) and possibly Mei-38 (the putative homologue of TPX2).

By using classical genetics, we have found that Cnn can recruit γ-tubulin complexes independently of Grip71 and the GCP4/5/4/6 core, meaning that it must be able to recruit γ-TuSCs. This is consistent with previous observations showing that γ-TuSCs could still be recruited to mitotic *Drosophila* centrosomes in S2 cells lacking the GCP4/5/4/6 core components (Vogt et al., 2006), although it was unknown at that time that this recruitment was dependent on Cnn. Our data here also shows that this occurs *in vivo*. The ability of Cnn to recruit γ-TuSCs is similar to its budding yeast homologues’ ability, where Grip71 and the GCP4/5/4/6 core are naturally absent. Consistent with this, we show that Cnn’s binding and recruitment of γ-tubulin complexes relies entirely on its highly conserved CM1 domain, which binds across the inter γ-TuSC interface in budding yeast complexes (Brilot et al., 2021). The binding of the CM1 domain in budding yeast stimulates the oligomerisation of γ-TuSCs into γ-TuRCs (Kollman et al., 2010; Lyon et al., 2016; Brilot et al., 2021; Lin et al., 2014; Gunzelmann et al., 2018a), but whether this is true of Cnn’s CM1 domain, or CM1 domains in other eukaryotes, remains to be determined. Consistent with this possibility, Cnn’s CM1 domain is more similar to Spc110’s, rather than Spc72’s, CM1 domain, which unlike Spc72 does not require the TOG domain protein Stu2 for efficient oligomerisation of γ-TuSCs. Moreover, Grip71 and the GCP4/5/4/6 core components are not essential in flies (Reschen et al., 2012; Vogt et al., 2006), nor in *Aspergillus* or *S. pombe* (Xiong and Oakley, 2009; Anders et al., 2006), suggesting that there must be ways to assemble ring-like templates in these organisms in the absence of the GCP4/5/4/6 core. We speculate that this “other way” is via CM1-mediated oligomerisation of γ-TuSCs (Figure 6). Nevertheless, Cnn and other CM1 domain proteins can also bind γ-TuRCs formed via the GCP4/5/4/6 core (Muroyama et al., 2016; Choi et al., 2010; Tovey et al., 2021; Wieczorek et al., 2020, 2019) and so it remains unclear whether Cnn recruits γ-TuSCs only in the absence of pre-formed γ-TuRCs.

In contrast to Cnn, Spd-2 (which does not contain a CM1 domain) requires Grip71 and the GCP4/5/4/6 core to recruit γ-tubulin complexes to mitotic centrosomes i.e. it is unable to bind and recruit γ-TuSCs directly or mediate their recruitment by another protein. Whether Spd-2 binds directly to preformed γ-TuRCs remains unclear. Grip71 associates with pre-formed γ-TuRCs in the cytosol and the human homologue of Grip71, NEDD1, has been reported to interact with the human homologue of Spd-2, CEP192 (Gomez-Ferreria et al., 2012a). Thus, Spd-2 might recruit γ-TuRCs via binding to Grip71, but since we show that Spd-2 can recruit γ-TuRCs in the absence of Grip71, it must also be able to recruit γ-TuRCs in a different way. Our data show that Grip163^GCP6^ is more important than Grip75^GCP4^ in this respect, because removing Grip71 and Grip75^GCP4^ does not completely abolish γ-tubulin accumulation. Perhaps, therefore, Spd-2 recruits γ-TuRCs via an interaction with Grip163^GCP6^ or Grip163^GCP6^, but not Grip75^GCP4^, is essential for the assembly of pre-formed γ-TuRCs necessary for Spd-2-mediated recruitment. This would be consistent with findings in human cells, where depletion of GCP6 is more disruptive to γ-TuRC assembly (Cota et al., 2017). So far, our attempts to identify direct interactions between Spd-2 and γ-TuRC components have failed, and so it is possible that an intermediary protein links Spd-2 to γ-TuRCs.

The finding that Cnn and Spd-2 recruit different types of γ-tubulin complexes to centrosomes fits well with recent observations that not all γ-TuRCs within a given species or cell type have the same protein composition. This was shown by analysing the γ-TuRC protein Mzt1 in *Drosophila*, *C. elegans*, fission yeast and *Aspergillus*, where Mzt1 is either not present or not necessary at certain MTOCs (Tovey et al., 2018; Gao et al., 2019; Huang et al., 2020; Sallee et al., 2018). For example, we have shown that *Drosophila* Mzt1 is expressed only in developing sperm cells and is required for γ-TuRC recruitment to basal bodies but not mitochondria (Tovey et al., 2018). Moreover, in mouse epithelial cells γ-TuRCs are bound and recruited either by CDK5RAP2 (Cnn homologue) or by NEDD1 (Grip71 homologue) and this influences the nucleation and anchoring ability of the γ-TuRCs (Muroyama et al., 2016). Whether other forms of γ-TuRCs also exist and how this affects their function remains to be explored.

In addition to revealing details of centrosomal recruitment of γ-tubulin complexes, we have also shown that microtubules can be nucleated in their absence. We’ve known for some time that microtubules are present within cells after depletion of γ-tubulin or other key γ-TuRC proteins (Hannak et al., 2002; Strome et al., 2001; Sampaio et al., 2001; Sunkel et al., 1995; Tsuchiya and Goshima, 2021; Sallee et al., 2018; Rogers et al., 2008; Nakaoka et al., 2015; Wang et al., 2015; Gunzelmann et al., 2018b) and that certain non-centrosomal MTOCs naturally lack γ-tubulin (Nashchekin et al., 2016; Yang and Wildonger, 2020; Zheng et al., 2020; Mukherjee et al., 2020; Kitamura et al., 2010). Mounting evidence, including our work here, suggests that the ch-Tog/XMAP215/Msps/Alp14/Stu2 TOG domain family of proteins (which have microtubule polymerase activity) and the TPX2 family of proteins (which have microtubule stabilization activity) are important for microtubule nucleation. Depletion of TOG domain proteins from *Xenopus* egg extracts, *Drosophila* S2 and fat body cells, fission yeast cells, and budding yeast cells, and depletion of TPX2 from *Xenopus* egg extracts, severely impairs microtubule nucleation or organisation (Popov et al., 2002; Thawani et al., 2018; Zheng et al., 2020; Groen et al., 2009; Flor-Parra et al., 2018; Gunzelmann et al., 2018b; Rogers et al., 2008). TOG domain and TPX2 proteins have been shown to work together with γ-TuRCs (or microtubule seed templates) to promote microtubule nucleation (Thawani et al., 2018; Flor-Parra et al., 2018; Gunzelmann et al., 2018b; Consolati et al., 2020; King et al., 2020; Wieczorek et al., 2015). Consistent with this, co-depletion of γ-tubulin and the *Drosophila* TOG domain protein Msps did not delay non-centrosomal microtubule regrowth after cooling compared to single depletions in interphase S2 cells (Rogers et al., 2008). Nevertheless, several studies, mainly *in vitro*, have shown that TOG and TPX2 proteins can also function independently of γ-TuRCs to promote microtubule nucleation (Roostalu et al., 2015; Woodruff et al., 2017; Schatz et al., 2003; Slep and Vale, 2007; Ghosh et al., 2013; Thawani et al., 2018; King et al., 2020; Zheng et al., 2020; Tsuchiya and Goshima, 2021). Our data suggest that, unlike from non-centrosomal sites in interphase S2 cells, Msps can promote γ-TuRC-independent microtubule nucleation from centrosomes in mitotic larval brain cells. This difference may reflect Msps having a high local concentration at centrosomes. This finding is similar to that of a recent study in human colon cancer cells showing that γ-tubulin depletion did not prevent microtubule nucleation from centrosomes and that this γ-TuRC-independent microtubule nucleation pathway depended on the Msps homologue ch-TOG (Tsuchiya and Goshima, 2021). It also supports the observation that *C. elegans* centrosome-like condensates nucleate microtubules with help from the TOG domain protein Zyg-9 (Woodruff et al., 2017). It is possible that Patronin is also involved in γ-TuRC-independent microtubule nucleation from centrosomes, but we were unable to test this. We note, however, that endogenously-tagged Patronin-GFP is not readily detectable at mitotic centrosomes in larval brain cells (unpublished observations). Interestingly, α/β-tubulin does not concentrate at mitotic centrosomes in flies (Figure S2), unlike in *C. elegans* where this can promote microtubule nucleation (Woodruff et al., 2017; Baumgart et al., 2019).

So why are γ-TuRCs required at all? While microtubules can be nucleated independently of γ-TuRCs, nucleation or microtubules growth appears to be more efficient when γ-TuRCs are present ((Tsuchiya and Goshima, 2021; Hannak et al., 2002); this study). Naturally occurring γ-TuRC-independent microtubule nucleation at specialised MTOCs, such as the nuclear envelope of *Drosophila* fat body cells (Zheng et al., 2020), may not require a high frequency of microtubule nucleation events, perhaps because they build their microtubule arrays over a relatively long period of time. During cell division, however, many microtubules must be generated rapidly, possibly creating a requirement for γ-TuRCs to provide efficient microtubule nucleation. Indeed, depleting γ-TuRCs delays spindle assembly and results in spindle and chromosome defects (Sunkel et al., 1995; Sampaio et al., 2001; Colombié et al., 2006; Vérollet et al., 2006). Nevertheless, centrosomes lacking γ-

TuRCs can organise similar, if not higher, numbers of microtubules as in controls (as observed in *cnn^cm1^,grip71,grip163* mutant cells). These microtubules are, however, less dynamic, being more cold-resistant and so perhaps going through round of depolymerisation/polymerisation less frequently. This reduced dynamicity may impact spindle assembly. γ-TuRCs may also be important to set microtubule protofilament number and define microtubule polarity, and studies have implicated γ-tubulin or γ-TuRC proteins in the control microtubule dynamics and of cell cycle progression, independent of their microtubule nucleation roles (Oakley et al., 2015; Bouissou et al., 2009).

In summary, our data highlight the robustness of centrosomes to nucleate microtubules. We have shown that centrosomes can recruit different forms of γ-tubulin complexes (γ-TuSCs and γ-TuRCs) via multiple pathways and that they can nucleate and organise microtubules in the absence of γ-tubulin complexes. This γ-TuRC-independent mode of microtubule nucleation relies on the TOG domain protein Msps. This multi-pathway redundancy helps explain why centrosomes are such dominant MTOCs during mitosis. A seemingly important finding is that microtubules nucleated by different mechanisms have different properties. This concept is similar to how the plus end dynamics of yeast microtubules are a function of where the microtubules were nucleated (Chen et al., 2019). These unexpected observations deserve further investigation.

## Supporting information

Cideo S1

## Acknowledgements

This research was supported by the Centre National de la Recherche Scientifique CNRS, Université Paris Cité, a Wellcome Trust and Royal Society Sir Henry Dale fellowship awarded to PTC [105653/Z/14/Z], by a Chaire d’excellence grant from the IdEx Université Paris Cité ANR-18-IDEX-0001 awarded to PTC, by an ATIP Avenir award funded by the Fondation Bettencourt Schueller awarded to PTC, by an Association pour la Recherche sur le Cancer grant (PJA 20181208148) awarded to AG, and by the CNRS. We thank Jordan Raff for sharing antibodies and fly lines. We thank Roger Karess for his invaluable input and critical reading of the manuscript. The work benefited from the Imaging Facility, Department of Zoology, University of Cambridge, supported by Matt Wayland and a Sir Isaac Newton Trust Research Grant (18.07ii(c)), and from the ImagoSeine at the IJM, Paris. For the purpose of Open Access, the author has applied a CC BY public copyright license to any Author Accepted Manuscript version arising from this submission.

## Author Contributions

ZZ produced fly stocks by combining alleles, performed cell imaging experiments, in particular establishing and executing the cooling-warming assays, and analysed data. IB contributed to the revision of the manuscript by generating fly lines, performing cell imaging experiments, analysing data, and providing feedback on the manuscript. CT designed and performed the immunoprecipitation experiment, including cloning and protein purification of the Cnn fragments, and provided feedback on the manuscript. EY contributed to establishing the fixed cell cooling-warming experiment. AG obtained funding to generate the EB1-GFP fly line via InDroso with coordination from FB. PTC obtained funding, designed the study, generated fly stocks, performed cell imaging experiments, analysed data, and wrote the manuscript.

## Methods

### Transgenic and endogenously modified *Drosophila* lines

The Jupiter-mCherry (Callan et al., 2010), GFP-PACT (Martinez-Campos et al., 2004) and RFP-PACT (Conduit et al., 2010) alleles have been described previously. To delete the CM1 domain from Cnn, we first designed a pCFD4 vector (Port et al., 2014) containing two guide RNAs with the following target sequences: AACTCGCCCTTGCCGTCACA and GTGATGAGAAATGGCTCGAG. This vector was injected into flies containing the attP2 landing site by Rainbow Transgenic Flies, Inc. Camarillo, CA 93012, USA. Male flies were then crossed to females expressing nos-cas9 (BL54591) and the resultant embryos were injected by the Department of Genetics Fly facility, University of Cambridge, UK, with a homology vector encoding 1kb on either side of the deletion region (R98 to D167, inclusive). The resulting F0 flies were crossed to balancer lines and their progeny were screened by PCR for the deletion.

The endogenously-tagged EB1-GFP line was made using CRISPR-based genome editing by inDroso, France. An SSSS-eGFP-3’UTR-LoxP-3xP3-dsRED-LoxP cassette was inserted and then the selection markers were excised. The guide RNA sequences were not communicated and the company has now closed.

### Mutant alleles, RNAi lines and fly stocks

For *spd-2* mutants, we used the *dspd-2^Z35711^*mutant allele, which carries an early stop codon resulting in a predicted 56aa protein. Homozygous *dspd-2^Z35711^* mutant flies lack detectable Spd-2 protein on western blots and so the allele is therefore considered to be a null mutant (Giansanti et al., 2008). In our stock collection, this allele no longer produces homozygous flies (which is common for mutant alleles kept as balanced stocks for many years), and so we combined *dspd-2^Z35711^*with a deficiency that deletes the entire *spd-2* gene (*dspd-2^Df(3L)st-j7^*). On western blots, there was no detectable Spd-2 protein in brain extracts from flies carrying the *dspd-2^Z35711^*/ *dspd-2^Df(3L)st-j7^* hemizygous mutations (Figure S6B). For *cnn* mutants, we combined the *cnn^f04547^* and *cnn^HK21^*mutant alleles. The *cnn^f04547^* allele carries a piggyBac insertion in the middle of the cnn gene and is predicted to disrupt long Cnn isoforms, including the centrosomal isoform (Cnn-C or Cnn-PA) (Lucas and Raff, 2007). This mutation is considered to be a null mutant for the long Cnn isoforms (Lucas and Raff, 2007; Conduit et al., 2014). The *cnn^HK21^* allele carries an early stop codon after Cnn-C’s Q78 (Vaizel-Ohayon and Schejter, 1999) and affects both long and short Cnn isoforms – it is considered to be a null mutant (Eisman et al., 2009; Chen et al., 2017a). On western blots, there was no detectable Cnn-C protein in brain extracts from flies carrying the *cnn^f04547^* / *cnn^HK21^* hemizygous mutations (Figure S6A). For Grip71, we used the *grip71^120^* mutant allele, which is a result of an imprecise p-element excision event that led to the removal of the entire *grip71* coding sequence except for the last 12bp; it is considered to be a null mutant (Reschen et al., 2012). We combined this with an allele carrying a deficiency that includes the entire *grip71* gene (*grip71^Df(2L)Exel6041^*). On western blots, there is no detectable Grip71 protein in *grip71^120^*/ *grip71^df6041^* hemizygous mutant brains (see blots on CRB website, which were performed by us). For Grip75^GCP4^, we used the *grip75^175^*mutant allele, which carries an early stop codon after Q291. Homozygous *grip75^175^* mutant flies lack detectable Grip75^GCP4^ protein on western blots and so the allele is therefore considered to be a null mutant (Schnorrer et al., 2002). We combined this with an allele carrying a deficiency that includes the entire *grip75^GCP4^* gene (*grip75^Df(2L)Exel7048^*). In the absence of a working antibody, we have not confirmed the expected absence of Grip75^GCP4^ protein in *grip75^175^* / *grip75^Df(2L)Exel7048^*hemizygous mutant flies on western blots. For Grip128^GCP5^, we used the UAS-controlled *grip128*-RNAi^V29074^ RNAi line, which is part of the VDRC’s GD collection, and drove its expression using the Insc-Gal4 driver (BL8751), which is expressed in larval neuroblasts and their progeny. In the absence of a working antibody, we have not confirmed the absence or reduction of Grip128^GCP5^ protein on western blots. RNAi was used for *grip128^GCP5^* as its position on the X chromosome made generating stocks with multiple alleles technically challenging. For Grip163^GCP6^, we used the *grip163^GE2708^*mutant allele, which carries a p-element insertion between amino acids 822 and 823 (total protein length is 1351aa) and behaves as a null or strong hypomorph mutant (Vérollet et al., 2006). We combined this with an allele carrying a deficiency that includes the entire *grip163^GCP6^* gene (*grip163^Df(3L)Exel6115^*). In the absence of a working antibody, we have not confirmed the absence or reduction of Grip163^GCP6^ protein in *grip163^GE2708^* / *grip163^Df(3L)Exel6115^* hemizygous mutant flies on western blots. For Msps, we used the *msps^p^* and *msps^MJ15^*mutant alleles. The *msps^p^* allele carries a p-element insertion within, or close to, the 5’ UTR of the *msps* gene and results in a strong reduction, but not elimination, of Msps protein on western blots (Cullen et al., 1999). The *msps^MJ15^* allele was generated by re-mobilisation of the p-element (the genetic consequence of which has not been defined) and also results in a strong reduction, but not elimination, of Msps protein on western blots (Cullen et al., 1999; Lee et al., 2001). For TACC, we used the *tacc^stella^* allele which contain a p-element insertion of unknown localisation but that results in no detectable TACC protein on western blots (Barros et al., 2005). For Mei-38, we used the UAS-controlled *mei-38*-RNAi^HMJ23752^ RNAi line, which is part of the NIG’s TRiP Valium 20 collection, and drove its expression using the Insc-Gal4 driver (BL8751). In the absence of a working antibody, we have not confirmed the absence or reduction of Mei-38 protein on western blots. RNAi was used for *mei-38* as its position on the X chromosome made generating stocks with multiple alleles technically challenging. Moreover, the only available mutant of *mei*-38 affects a neighbouring gene.

For examining the behaviour of MTs in living larval brain cells, we analysed brains expressing 2 copies of Ubq-GFP-PACT and 2 copies of Ubq-Jupiter-mCherry in either a WT, a *cnn,grip71,grip163^GCP6^*mutant, or a *cnn*^Δ*CM1*^*,grip71,grip163^GCP6^*background. For examining the behaviour of EB1-GFP in living larval brain cells, we analysed brains expressing 2 copies of EB1-GFP in either a WT or a *cnn,grip71,grip163^GCP6^* mutant background.

### Antibodies

For immunofluorescence analysis, we used the following antibodies: mouse anti-γ-tubulin monoclonal (1:500; GTU88, Sigma), mouse anti-α-tubulin monoclonal (1:1000; DM1α, Sigma), rabbit anti-α-tubulin monoclonal (1:500; AB52866, Abcam), anti-PhosphoHistoneH3 monoclonal (mouse, 1:2000, Abcam or rabbit, 1:500, Cell Signalling Technology), Guinea pig anti-Asl polyclonal (1:500; Gift from Jordan Raff), rabbit anti-DSpd-2 polyclonal (1:500) (Dix and Raff, 2007). Secondary antibodies were from Molecular Probes (Invitrogen): Alexa Fluor 488, 568, and 647 (all used at 1:1000). For western blotting we used mouse anti-γ-tubulin monoclonal (1:250; GTU88, Sigma), rabbit anti-MBP polyclonal (1:1000; gift from Jordan Raff), rabbit anti-Cnn polyclonal (1:1000; gift from Jordan Raff), rabbit anti-Spd-2 polyclonal (1:500; gift from Jordan Raff), and rabbit anti-Grip71 polyclonal (1:250; CRB #2005268).

### Fixed brain analysis

For the analysis of centrosomal fluorescence levels of γ-tubulin or other PCM components, 3^rd^ instar larval brains were processed as described previously (Conduit et al., 2014). Briefly, dissected larval brains were incubated in 100μM colchicine in Schneider’s medium for 1h at 25°C to depolymerise microtubules. This prevents centrosomes in *cnn* mutants from “rocketing” and transiently losing their PCM (Lucas and Raff, 2007), allowing a more accurate quantification of PCM recruitment (Conduit et al., 2014). Brains were fixed in 4% paraformaldehyde containing 100mM PIPES, 1mM MgSO_4_, and 2mM EGTA pH 6.95 for 20 minutes at room temperature, washed in PBS and then 45% and 60% acetic acid, squashed under a coverslip, post-fixed in methanol, washed in PBT, and then stained with the appropriate antibodies. Images were collected on either a Leica SP5 point scanning upright confocal system run by LAS AF software using a 63X 1.3NA glycerol objective (Leica 1156194), or a Zeiss Axio Observer.Z1 inverted CSU-X1 Yokogowa spinning disk system with 2 ORCA Fusion camera (Hamamatsu) run by Zeiss Zen2 acquisition software using a 60X 1.4NA oil immersion lens (Zeiss), or a Zeiss LSM700 confocal microscope. At least 5 images containing multiple cells in both mitosis (as shown by positive Phospho-Histone H3 staining) and interphase were collected for each brain. Each data point on a graph represents the average signal from all the centrosomes quantified in a single brain. Typically, between 30 and 50 centrosomes were analysed per cell cycle stage (interphase or mitosis) per brain.

For assessing the ability of centrosomes to organise microtubules during prophase, 3^rd^ instar larval brains were treated and imaged as above except that the colchicine incubation step was omitted. A prophase cell was scored as positive when at least one centrosome had an associated α-tubulin signal. For measuring the distance of centrosomes from the spindle pole during prometaphase, measurements were made between the centre of Asl signal (centrosome) and the spindle pole (centre of the α-tubulin signal at the spindle pole).

### Fixed microtubule re-growth assay

3^rd^ instar larval brains of the appropriate genotype were dissected and incubated on ice in Schneider’s medium for 40 minutes. Empirical tests showed that a 40-minute incubation was necessary to efficiently depolymerise centrosomal microtubules. Larval brains were then either rapidly fixed on ice in 16% paraformaldehyde containing 100mM PIPES, 1mM MgSO_4_, and 2mM EGTA pH 6.95 for 5 minutes (T0 brains), or were quickly transferred to room temperature for 30 seconds and then rapidly fixed at room temperature. Subsequently, the brains were processed as above.

### Live analysis of microtubule and EB1-GFP comets during cooling warming cycles

A CherryTemp device from CherryBiotech was used to modulate the temperature of larval brain cells. 3^rd^ instar larval brains were dissected and semi-squashed between a coverslip and the CherryTemp thermalisation chip in Schneider’s medium. The *cnn,grip71,grip163* mutant samples and their respective controls were imaged on a Leica DM IL LED inverted microscope controlled by μManager software and coupled to a RetigaR1 monochrome camera (QImaging) and a CoolLED pE-300 Ultra light source using a 63X 1.3NA oil objective (Leica 11506384). The *cnn*^Δ*CM1*^*,grip71,grip163* mutant samples and their respective controls were imaged on a Leica DMi8 inverted microscope controlled by μManager software and coupled to a BSI Prime Express monochrome camera (QImaging) and a CoolLED pE-300 Ultra light source using a 63X 1.3NA oil objective (Leica 11506384). The temperature was changed from 20°C to 5°C and back to 20°C for microtubule depolymerisation and repolymerisation, respectively. Temperature changes induce movements in the glass and the focus was manually adjusted to keep as many frames in focus as possible during the temperature shifts. For Jupiter-mCherry Videos, Z-stacks with gaps of 500nm were acquired every 6 seconds; for EB1-GFP Videos, Z-stacks with gaps of 300nm were acquired every second. For the quantification of Jupiter-mCherry in the *cnn,grip71,grip163* mutant experiment, 12 and 10 centrosomes from 7 and 10 control and *cnn,grip71,grip163* mutant cells were analysed, respectively. For the quantification of Jupiter-mCherry in the *cnn*^Δ*CM1*^*,grip71,grip163* mutant experiment, 12 and 11 centrosomes from 4 and 4 control and *cnn*^Δ*CM1*^*,grip71,grip163* mutant cells were analysed, respectively. GraphPad Prism was used to generate the one-phase exponential decay models and exponential plateau models that are fitted to the depolymerisation and nucleation/regrowth phases, respectively.

### Image analysis and statistics

All images were processed in Fiji (Image J). Each Z-stack image was reconstructed by maximum intensity Z-axis projection. PCM or microtubule levels at centrosomes were calculated by measuring the total fluorescence in a boxed or circular region around the centrosome and subtracting the local cytoplasmic background fluorescence. GraphPad Prism was used for statistical analysis. Details of N numbers, the statistical tests and models used can be found in the figure legends.

### Recombinant protein cloning, expression and purification

The Cnn-C-N^T27E,S186D^ fragment comprises amino acids 1-255 of Cnn-C and was generated previously (Tovey et al., 2021). We generated the Cnn-C-N^T27E,R101Q,E102A,F115A,S186D^ fragment in a similar manner. Briefly, a pDEST-HisMBP (Addgene, #11085) vector containing aa1-255 of Cnn was digested with KpnI and SspI and a complementary fragment containing the point mutations was cloned into the cut vector using HiFi technology (NEB). The complementary fragment was generated by GENEWIZ. The vector was transformed into BL21-DE3 cells and the protein purified using gravity flow amylose resin (New England Biolabs) affinity chromatography. Peak elution fractions were diluted 1:1 with glycerol and stored at - 20°C.

### Immunoprecipitation and western blotting

Immunoprecipitation was carried out as previously (Tovey et al., 2021). Briefly, 1g/ml of wild-type embryos were homogenised in homogenisation buffer containing 50 mM HEPES, pH7.6, 1mM MgCl_2_, 1 mM EGTA, 50 mM KCl supplemented with PMSF 1:100, Protease Inhibitor Cocktail (1:100, Sigma Aldrich) and DTT (1M, 1:1000). Extracts were clarified by centrifugation twice for 15 minutes at 16,000 rcf at 4°C and 100 μl embryo extract was rotated at 4°C overnight with 30 μl magnetic ProteinA dynabeads (Life Technologies) coupled with anti-MBP antibodies (gift from Jordan Raff) and MBP-Cnn fragments. Beads were washed 5 times for 1 min each in PBS + 0.1% triton (PBST), boiled in 50 μl 2x sample buffer (BioRad), and separated from the eluted IP sample using a magnet. Samples were analysed by electrophoresis and western blotting as described previously (Tovey et al., 2021). Membranes were imaged using a BioRad ChemiDoc.

### Summary of Supplementary material

The paper includes 6 Supplemental Figures and 4 Supplemental Videos.

---

**Figure S1**

**Centrosomes from *cnn***^Δ***CM1***^ **mutants accumulate slightly more** γ**-tubulin than centrosomes from *cnn* null mutants. (A)** Fluorescence images of mitotic *Drosophila* brain cells from either *cnn* or *cnn*^Δ*CM1*^ mutant third instar larvae immunostained for γ-tubulin (green), mitotic DNA (blue), and Asl (centrioles, magenta). Note how the γ-tubulin signal in *cnn*^Δ*CM1*^ mutant cells is also offset from the Asl signal, indicating that removing the CM1 domain affects the Cnn’s ability to form a proper centrosomal scaffold. Scale bars are 5μm. **(B)** Graph showing average fluorescence intensities of interphase (blue dots) and mitotic (black dots) centrosomes from either *cnn* or *cnn*^Δ*CM1*^ mutant brains (as indicated below). Each data-point represents the average centrosome value from one brain. Mean and SEM are indicated. Brains from the different genotypes were paired on slides (one slide per pair) allowing a paired t-test to compare the mean values between mitotic centrosomes. Note that γ-tubulin accumulation at mitotic centrosomes is only slightly higher in *cnn*^Δ*CM1*^ mutant cells, indicating that either the Cnn-dependent pool of Spd-2 is not an efficient recruiter of γ-TuRCs or that recruitment of the Cnn-dependent pool of Spd-2 is perturbed in *cnn*^Δ*CM1*^ mutant cells, or both.

**Figure S2**

**Drosophila centrosomes do not concentrate** α**/**β**-tubulin.** Fluorescence images of mitotic *Drosophila* brain cells from wild-type third instar larval brains that had been cooled for 40 minutes on ice immunostained for alpha-tubulin (microtubules, green), mitotic DNA (blue), and Asl (centrioles, magenta). Note how there is no tubulin signal at centrosomes that is above cytosolic levels.

**Figure S3**

**Plots of individual centrosome Jup-mCherry values during cooling-warming experiments. (A,B)** Graphs plotting the centrosomal signal (after subtraction of cytosolic signal) of Jupiter-mCherry within living *Drosophila* control (D) and *cnn,grip71,grip163* mutant (E) third instar larval brain cells as they were cooled to 5°C for ∼3 minutes and then rapidly warmed to 20°C. Time in seconds relative to the initiation of warming (0s) is indicated. Note how the centrosomal Jupiter-mCherry signal does not always reach cytosolic levels (i.e. 0), indicating that microtubules were not fully depolymerised from all centrosomes, but note also that even when the Jupiter-mCherry signal did reach cytosolic levels there was still a rapid increase after warming, indicating that the increase in signal after warming is not simply due to regrowth of partially depolymerised microtubules.

**Figure S4**

**The distribution of cells with and without centrosomes and with and without re-formed spindles is similar in *cnn,grip71,grip163* mutant and *cnn,grip71,grip163,msps* mutant cells after 30 seconds of warming post cooling.** Parts of a whole graphs show the proportion of cells that either contain centrosomes or do not, and that have either formed a spindle or have not, in cells fixed and immunostained for alpha-tubulin (microtubules, green), mitotic DNA (blue), and Asl (centrioles, magenta) after 30 seconds of warming post cooling from either *cnn,grip71,grip163* or *cnn,grip71,grip163,msps* mutants, as indicated.

**Figure S5**

**Spindles form less robustly in cells depleted of Msps in addition to Cnn, Grip71 and Grip163^GCP6^. (A-C)** Parts of a whole graphs (A,C) and images (B) represent analyses of mitotic spindle quality in cells fixed and immunostained for alpha-tubulin (microtubules, green), mitotic DNA (blue), and Asl (centrioles, magenta) after 30 seconds of warming post cooling from either wild-type, *cnn,grip71,grip163* or *cnn,grip71,grip163,msps* mutants, as indicated. Only cells that had centrosomes and that had already formed a spindle were analysed in (A,B) and only cells lacking centrosomes but that had formed a spindle were analysed in (C). Note that wild-type cells always contained centrosomes and so were not analysed in (C). Note also how spindle quality is frequently low in *cnn,grip71,grip163,msps* mutants (A,B), but the difference in spindle quality between mutant types is not apparent in cells lacking centrosomes (C). Note also that some cells lacking Cnn have abnormal numbers of centrosomes due to centrosome segregation problems during cell division (Conduit et al., 2010).

**Figure S6**

**Brains from flies carrying mutant alleles for *cnn* or *spd-2* display no observable Cnn or Spd-2 protein on western blots.** Western blots of larval brain samples from wild-type and cnn,spd-2,pUbq-RFP-Cnn flies probed with anti-Cnn (**A**) and anti-Spd-2 (**B**). Note how anti-Cnn recognises endogenous Cnn in the wild-type sample and the larger exogenous pUbq-RFP-Cnn in the cnn,spd-2,pUbq-RFP-Cnn mutant sample, but that no endogenous Cnn is detected in the cnn,spd-2,pUbq-RFP-Cnn mutant sample. Note also how anti-Spd-2 recognises endogenous Spd-2 in the wild-type sample but not in the cnn,spd-2,pUbq-RFP-Cnn mutant sample.

## Supplementary Videos

**Video 1**

**Microtubule depolymerisation and regrowth at centrosomes in control cells.** Video showing the Jupiter-mCherry signal (marking microtubules) within a living control cell during a cooling-warming experiment. Cooling to 5°C begins at −174s and warming to 20°C begins at 0s. Note how the Jupiter-mCherry signal decreases gradually during cooling and then recovers immediately at the two centrosomes during warming.

**Video 2**

**Microtubule depolymerisation and regrowth at centrosomes in *cnn,grip71,grip163* mutant cells.** Video showing the Jupiter-mCherry signal (marking microtubules) within a living *cnn,grip71,grip163 mutant* cell during a cooling-warming experiment. Cooling to 5°C begins at −186s and warming to 20°C begins at 0s. Only one centrosome is present (spindle pole on the left). Note how the Jupiter-mCherry signal decreases gradually during cooling and then recovers immediately at the centrosome during warming, but that this recovery is slower and less intense than in control cells.

**Video 3**

**Behaviour of EB1-GFP comets at centrosomes during a cooling-warming cycle in control cells.** Video showing the EB1-GFP signal (marking growing microtubule ends) within a living control cell during a cooling-warming experiment. Cooling to 5°C begins at −54s and warming to 20°C begins at 0s. Note how the EB1-GFP signal disappears immediately on cooling and then dramatically reappears and spreads outwards from the two centrosomes during warming.

**Video 4**

**Behaviour of EB1-GFP comets at centrosomes during a cooling-warming cycle in *cnn,grip71,grip163 mutant* cells.** Video showing the EB1-GFP signal (marking growing microtubule ends) within a **living *cnn,grip71,grip163* mutant** cell during a cooling-warming experiment. Cooling to 5°C begins at −276s and warming to 20°C begins at 0s. Note how the EB1-GFP signal does not disappear immediately on cooling, unlike in control cells. The signal does disappear fully prior to warming and then reappears from the centrosomes and chromosomal regions during warming, spreading outwards. Note also that the centrosomes do not remain in focus throughout the movie. The centrosome at the spindle pole in the lower half of the cell is in focus throughout most of the movie, but this centrosome is out of focus for ∼30s after warming due to fluctuations in the cover glass during the temperature change.

## Notes

### Competing Interest Statement

The authors have declared no competing interest.

### Summary of Updates

we have added a cnn mutant only control to Figure 2B, we have compared g-tubulin recruitment between cnn and cnn∆CM1 mutants (new Figure S1), we have shown that g-tubulin does not accumulate at mitotic centrosomes in cnn∆CM1,grip71, grip163 mutants (new Figure 4A,B), and we have quantified the dynamics of the centrosomal microtubules in these mutants (new Figure 4C-G). We have also adjusted the text to reduce the character count.

