## Supplementary figures and images for "Multifaceted modes of γ-tubulin complex recruitment and microtubule nucleation at mitotic centrosomes"

### Figure S1

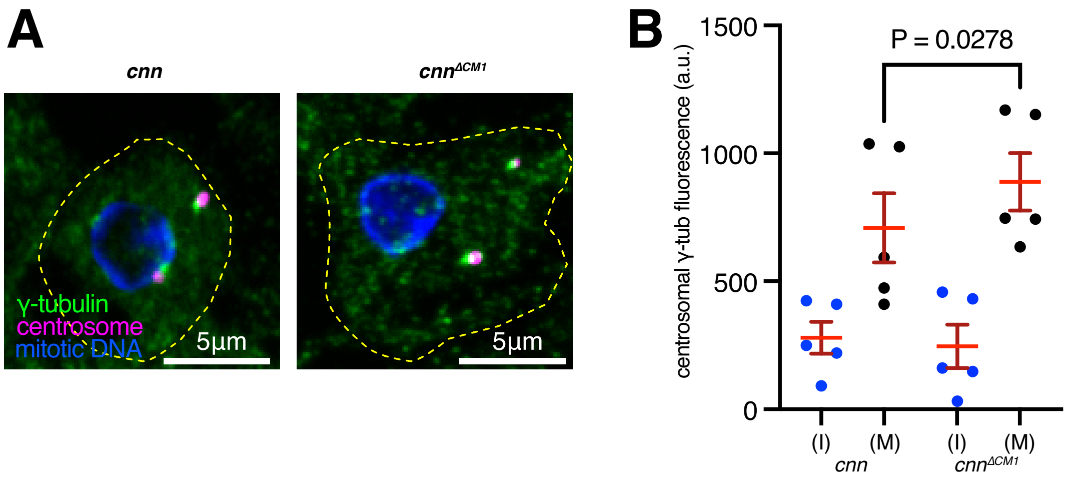

### Figure S2

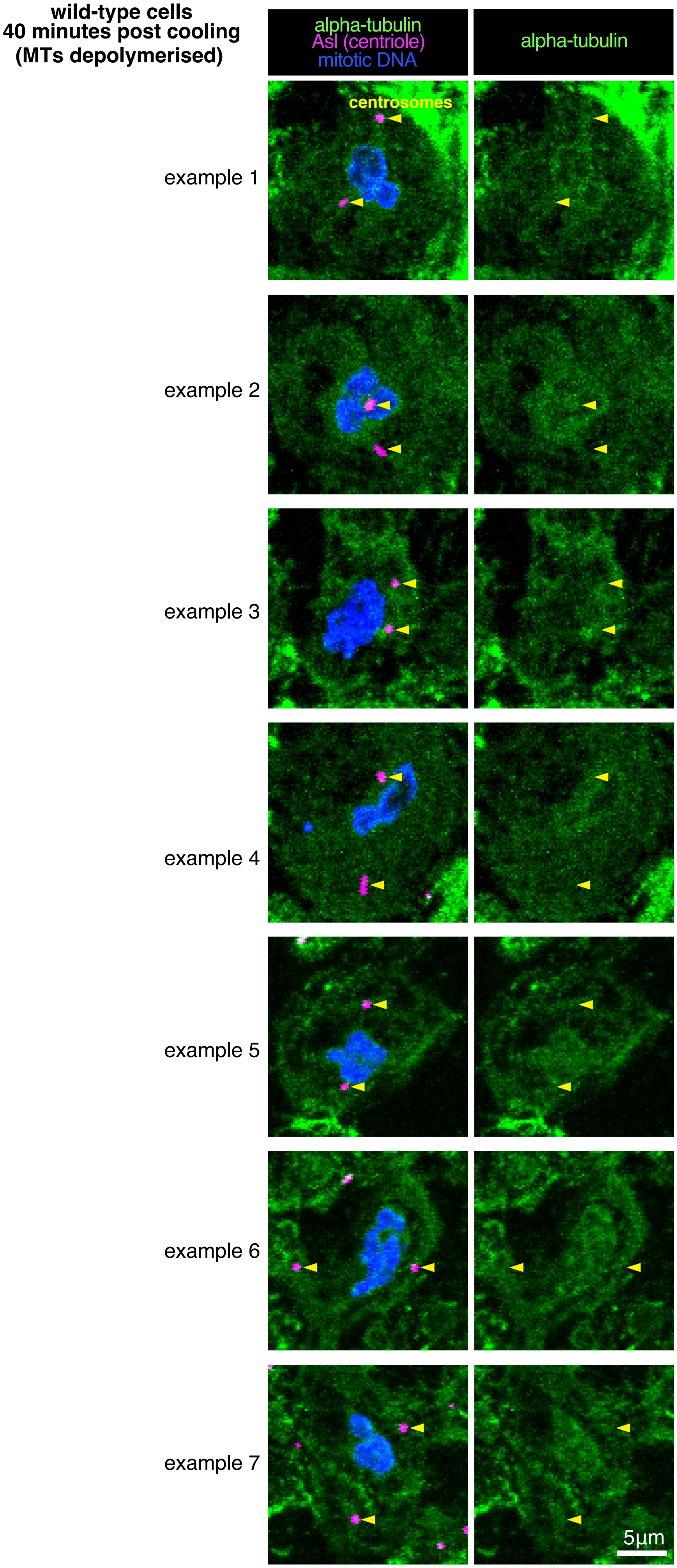

### Figure S3

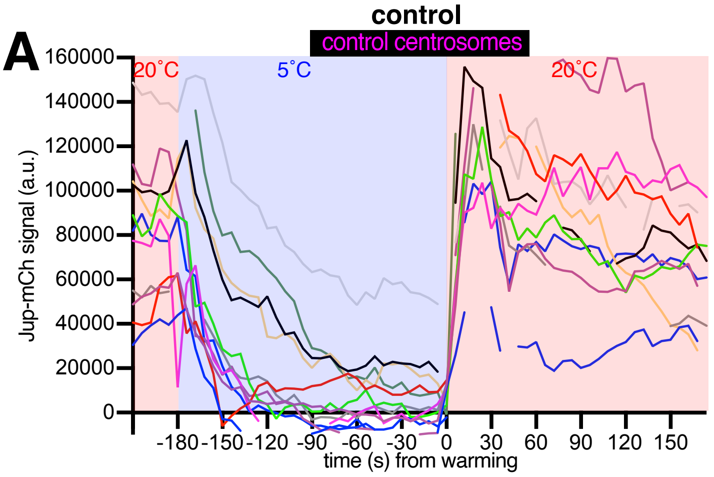

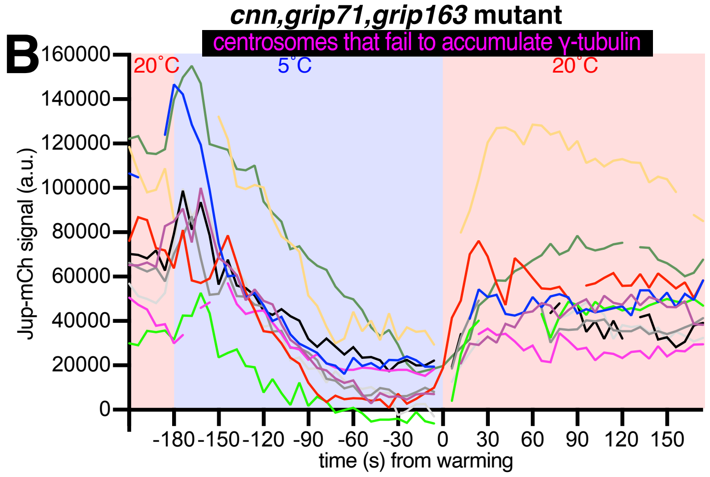

### Figure S4

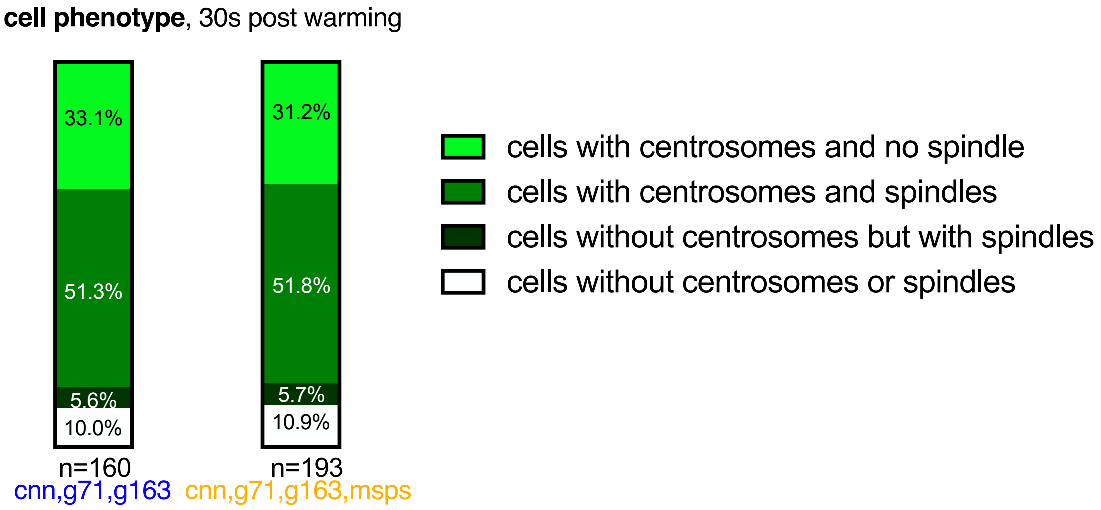

### Figure S5

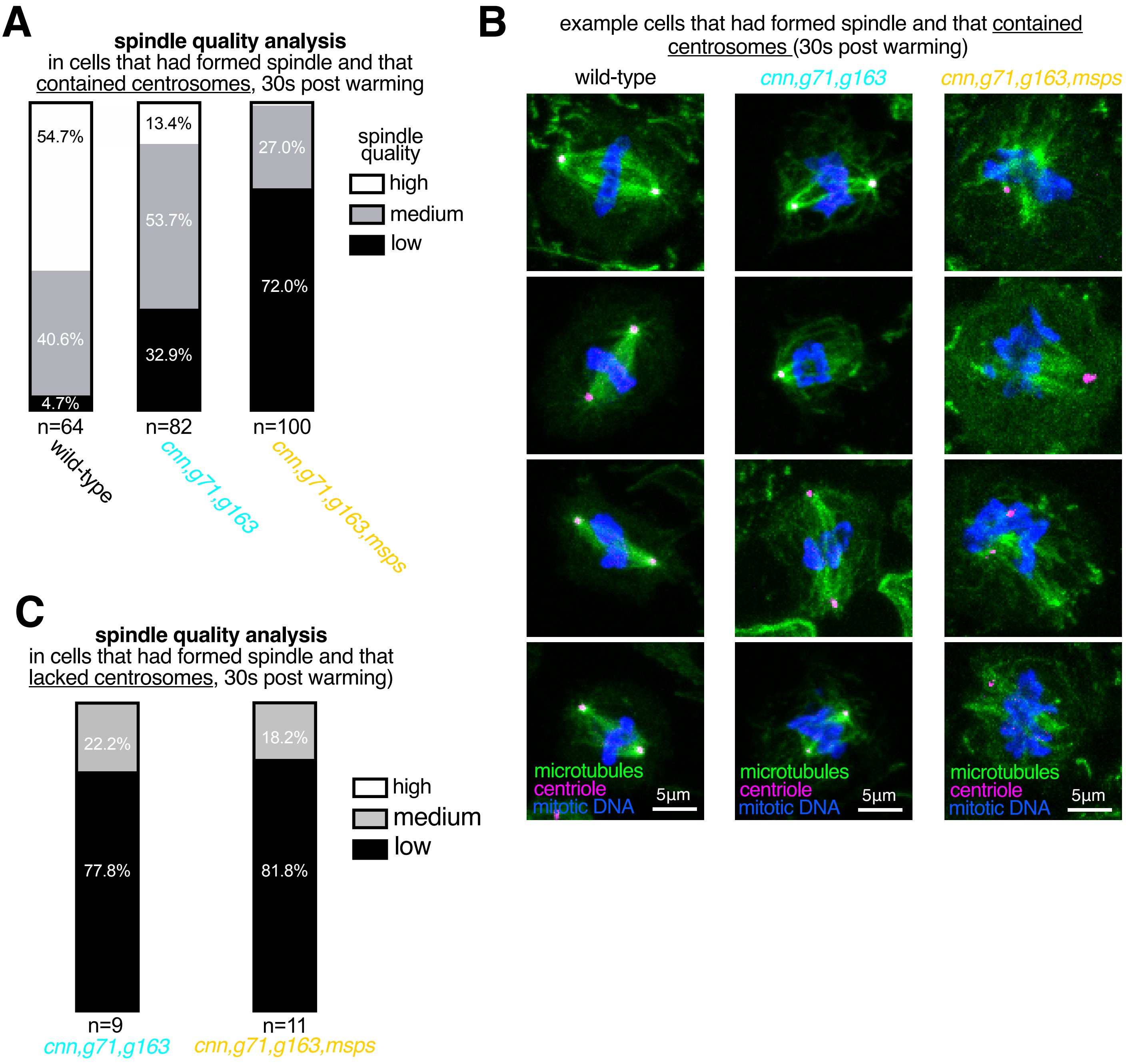

### Figure S6

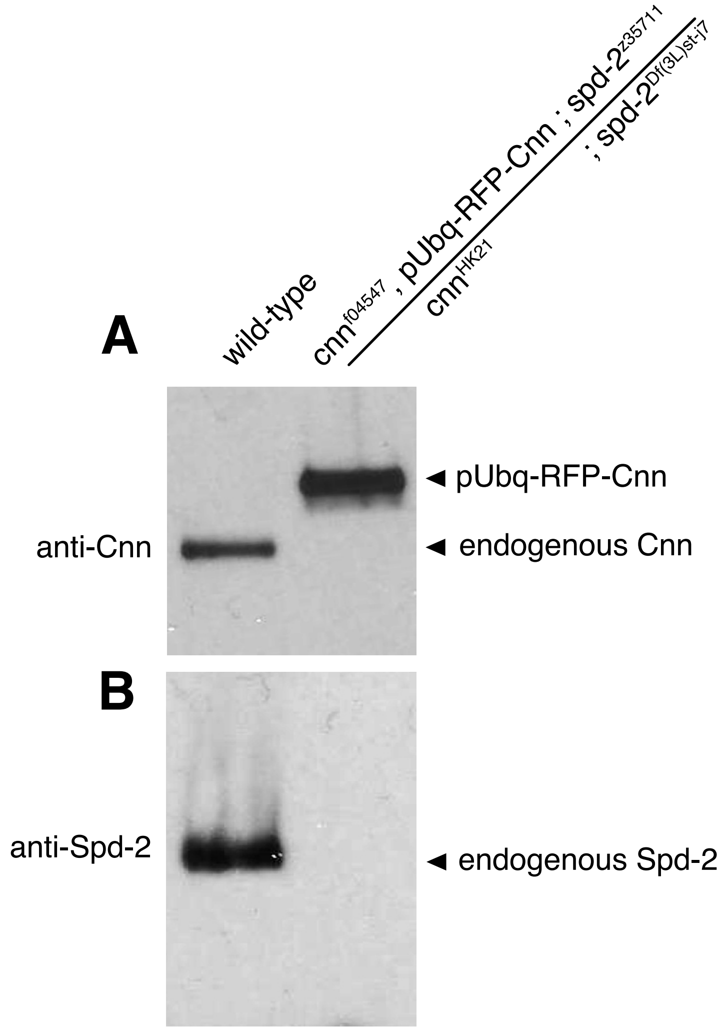
